# CLEAR-IT: Contrastive Learning to Capture the Immune Composition of Tumor Microenvironments

**DOI:** 10.1101/2024.08.20.608738

**Authors:** Daniel Spengler, Serafim Korovin, Kirti Prakash, Peter Bankhead, Reno Debets, Carlas Smith, Hayri E. Balcioglu

**Affiliations:** Delft Center for Systems and Control, Delft University of Technology, Delft, the Netherlands; Centre for Genomic & Experimental Medicine, Institute of Genetics and Cancer, The University of Edinburgh, Edinburgh, United Kingdom; Edinburgh Pathology and CRUK Scotland Centre, Institute of Genetics and Cancer, The University of Edinburgh, Edinburgh, United Kingdom; Laboratory of Tumor Immunology, Department of Medical Oncology, Erasmus MC Cancer Institute, Erasmus Medical Center, Rotterdam, the Netherlands

**Keywords:** Contrastive Learning, Self-supervised Learning, Cell Phenotyping, Tumor Microenvironment, Multiplexed Imaging, Sparse Labeling

## Abstract

Accurate phenotyping of cells in the tumor microenvironment is crucial for understanding cancer biology and developing effective therapies. However, current methods require precise cell segmentations and struggle to generalize across different imaging modalities, limiting their utility in digital pathology. Here, we show that Contrastive Learning Enabled Accurate Registration of Immune and Tumor Cells (CLEAR-IT) overcomes these limitations, providing a robust and versatile tool for cell phenotyping. CLEAR-IT accurately phenotypes cells comparable to state-of-the-art methods, generalizes across multiplex imaging modalities, maintains high performance even with limited number of labels, and enables the extraction of prognostic markers. Additionally, CLEAR-IT can be combined with existing methods to boost their performance, whereas its lack of need for precise cell segmentations significantly reduces training efforts. This method enhances the robustness and efficiency of digital pathology workflows, making it a valuable tool for cancer research and diagnostics.

---

Recent advances in multiplexed proteomics have underscored the critical role of tumor microenvironment (TME) spatial composition in cancer prognosis and therapy development [1–4]. As manual characterization of increasing amounts of data becomes infeasible, machine learning algorithms for TME characterization are being developed [5–8]. However, these approaches often require precise cell segmentation and extensive expert annotations, limiting their usability (**Fig. 1a**). Contrastive self-supervised learning can maintain high performance with fewer labels in image classification [9], and has been applied successfully to tissue segmentation and histopathological image classification [10], but not yet to multiplex images for the purpose of single-cell phenotyping.

**Fig. 1.**
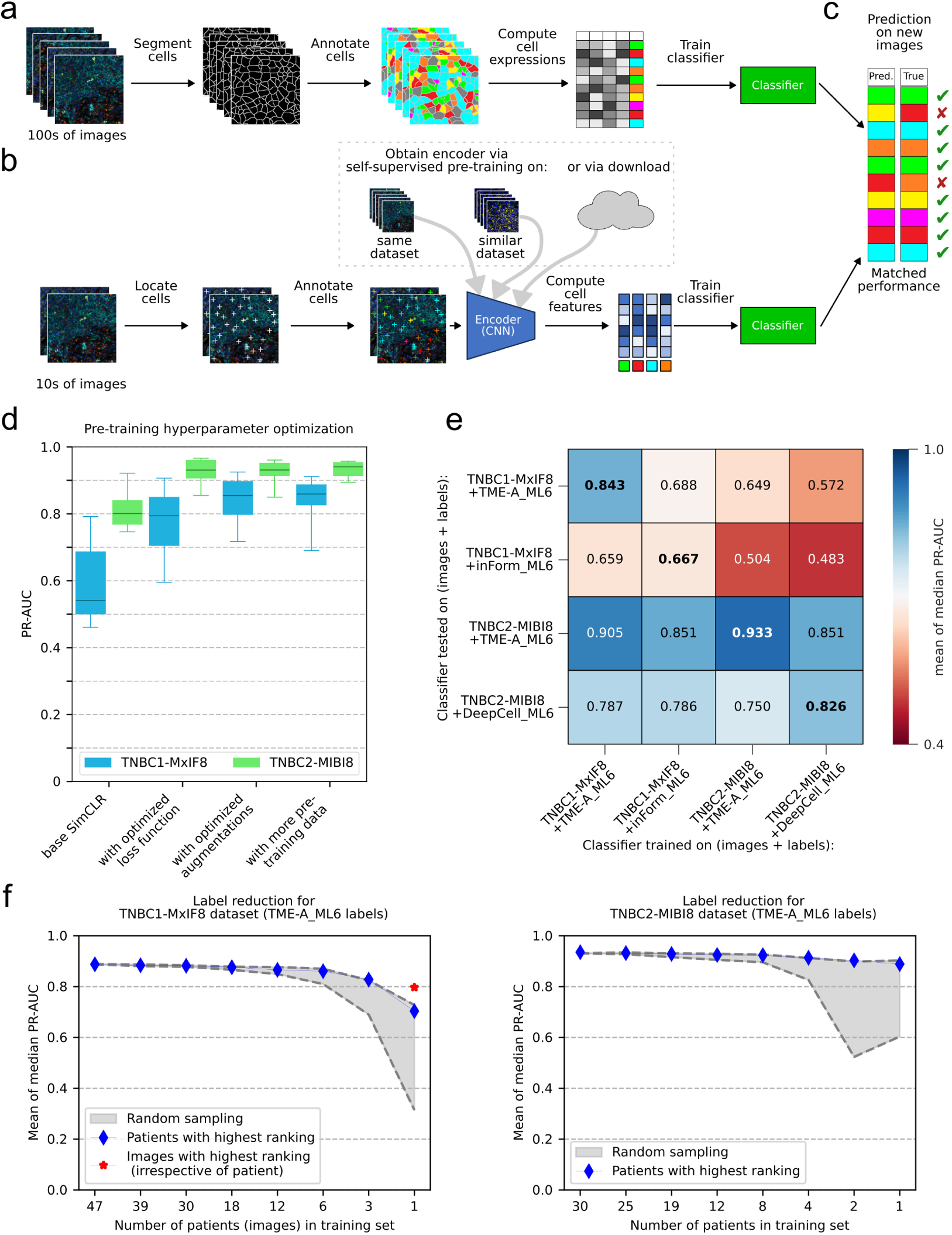
CLEAR-IT pipeline and performance. **a)** Conventional training pipelines for cell phenotyping are trained on cell expressions which require large amounts of segmentation ground truth. **b)** Use of feature encodings generated by CLEAR-IT; convolutional neural network (CNN) encoder trained self-supervised without annotations; reduces hands-on-time needed by increasing label efficiency and using only cell locations. **c)** Matched performance is achieved between benchmark and CLEAR-IT. **d)** PR-AUC box-and-whisker plots showing CLEAR-IT-SLP gain of performance for the TNBC1-MxIF8 and TNBC2-MIBI8 datasets through optimization of the pre-training hyperparameters for the CLEAR-IT encoder. **e)** Heatmap showing the cross-dataset PR-AUC scores of CLEAR-IT-SLP for TNBC1-MxIF8 and TNBC2-MIBI8 datasets. **f)** CLEAR-IT-SLP PR-AUC scores of supervised trained SLPs as a function of number of patients labels originated from for TNBC1-MxIF8 (left) and TNBC2-MIBI8 (right) datasets; shaded gray regions: performance range of 10 random patients; blue diamonds: patients with the highest ranking; red star: labels from 21 images with the highest ranking irrespective of patient of origin.

Contrastive Learning Enabled Accurate Registration of Immune and Tumor cells (CLEAR-IT) is a contrastive learning-based classifier designed to identify cell types in multiplexed tissue images using sparsely labeled data and only cell locations without the need for accurate segmentation (**Fig. 1b,c**). It utilizes the “Simple Framework for Contrastive Learning of Visual Representations” (SimCLR) algorithm for self-supervised learning [10]. For CLEAR-IT optimization, the following five steps are used; (i) image patches around cell centroids are made; (ii) an encoder is pre-trained on single-channel image patches extracted from multiplexed datasets without annotation; (iii) the encoder outputs are concatenated to obtain multi-channel representation; (iv) a single-layer perceptron (SLP) is trained with annotated cell phenotype data to obtain cell phenotype probabilities; (v) which is then compared with ground-truth annotations to quantify precision-recall area-under-the-curve (PR-AUC) to evaluate the performance (**Methods**, **Fig. S1**). Here, steps i–iii represent the CLEAR-IT approach, and PR-AUC quantification of CLEAR-IT-SLP allows direct evaluation of the CLEAR-IT performance on the task of cell phenotyping.

To ensure high performance, CLEAR-IT was optimized through iterative adjustments to the pre-training loss function, augmentations, and training data, evaluating the cell phenotyping performance of CLEAR-IT-SLP (**Fig. S1c**), separately in two different triple negative breast cancer (TNBC) cohorts (**Supplementary Information**). These cohorts contained 1010 8-channel multiplex immunofluorescence images (TNBC1-MxIF8) [1] or 41 44-channel multiplexed ion beam imaging by time-of-flight images (TNBC2-MIBI44) [4], latter reduced to 8-channel images for direct comparison with TNBC1-MxIF8 (TNBC2-MIBI8). For ground truth, multi-label annotations with 6 different class labels generated with TME-Analyzer [11] (TME-A ML6) were used (**Supplementary Information**). For both datasets, final PR-AUC was *>* 0.8, with highest performance gain of *>*30 % obtained via optimization of the loss function and augmentation optimizations resulting in an additional ∼15 % gain, whereas increasing the training data size, also with data from other sources, did not result in additional performance gain (**Fig. 1d**, **Fig. S2**–**Fig. S11**). Evaluating the performances and cross-performances of different CLEAR-IT-SLPs, obtained across imaging and analysis platforms, demonstrated highest performance for “self-classifiers” (**Fig. 1e**, main diagonal), where training was performed on the same dataset as the testing. Here, also published multi-class annotations [1, 4] were used to generate additional multi-label annotations (inForm ML6 for TNBC1-MxIF8 and DeepCell ML6 for TNBC2-MIBI8, **Supplementary Information**) to compare different analysis platforms. For the cross-performances, training on TNBC1-MxIF8 resulted in higher performance than training on TNBC2-MIBI8, and classifiers trained with TME ML6 annotations outperformed other annotations, resulting in the highest overall performance by TNBC1-MxIF8 + TME-A ML6, where performance loss compared to “self-classifiers” was 0 % to 5 % (**Fig. 1e**, **Fig. S12**), demonstrating that CLEAR-IT generalizes well to different imaging and analysis platforms.

Testing the resilience of our approach to label reduction in “self-classifiers” demonstrated high performance down to 10 % of our training data, corresponding to 0.1 % of all available data, with the exception of TNBC2-MIBI8 + DeepCell ML6 where performance dropped significantly below 20 % of training data (1 % of available data, **Fig. S13**). Reduction of labels in the form of number of patients, which represents a more realistic scenario, showed that with 10 % of available patients (TNBC1-MxIF8: 6 patients; TNBC2-MIBI8: 4 patients) less than 10 % reduction in performance was achieved for all cases (**Fig. S14**). With high confidence true annotations corresponding to high intensity signals at the center of the image (**Fig. E1**, **Fig. S15** and **Fig. S16**) and, despite large variations in the performance of CLEAR-IT-SLP when the SLP was trained with labels from one patient, high overall signal intensity distribution relating to high performance (**Fig. E2**, **Table S1**), we concluded that signal intensity was central to CLEAR-IT performance. We therefore developed a ranking scheme based on image signal intensity standard deviation (**Fig. E3a**, **Methods**), obtaining correlation between patient ranking and the CLEAR-IT-SLP performance, when the SLP was trained using data solely from one patient (**Fig. E3b,c**, *r* = 0.49 for TNBC1-MxIF8 and *r* = 0.9 for TNBC2-MIBI8, *p <* 0.001 for both). Notably, the performance with the best ranking patient was 18.6 % and 4.6 % lower than a full training cohort, for TNBC1-MxIF8 and TNBC2-MIBI8, respectively (**Fig. 1f**, blue diamonds). Here, the best ranking patient in the TNBC1-MxIF8 dataset had 21 images, compared to 1 image in the TNBC2-MIBI8 dataset, and training the SLP in TNBC1-MxIF8 dataset with 21 highest ranking images, irrespective of patient of origin, resulted in a mere 10.1 % reduction in performance over the training with 761 images from 47 patients (**Fig. 1f**, red asterisk). This highlights the image selection for data annotation as a crucial step that can significantly boost performance. This, together with high performance obtained with reduced labeled data with CLEAR-IT, can significantly reduce the expert annotation needed with minimal performance loss.

Having established high label efficiency through image ranking, we next investigated if clinically relevant parameters can be extracted in this scenario. Using labels generated by CLEAR-IT-SLP trained with minimal labels (TNBC1-MxIF8: **Fig. 1f**, left, red asterisk; TNBC2-MIBI8: **Fig. 1f**, right, 1 patient blue diamond) and our recently published methodology [11], we were able to obtain survival classifiers that split the cohorts into two differential overall survival groups (**Fig. 2a,b**, top). These classifiers were cross-validated (**Fig. 2a,b**, bottom), where classifier training was blind to the information from the validation cohort. This demonstrated that CLEAR-IT, even at highly reduced label presence, can prognosticate TNBC patients. Furthermore, when investigating the parameters that are involved in survival classifiers, we observed that the parameter ranking generated by CLEAR-IT-SLP were in good agreement with the published ranking [11] (**Fig. 2c**). This showed that CLEAR-IT is able to extract clinically relevant TME biology, with minimal user input.

**Fig. 2.**
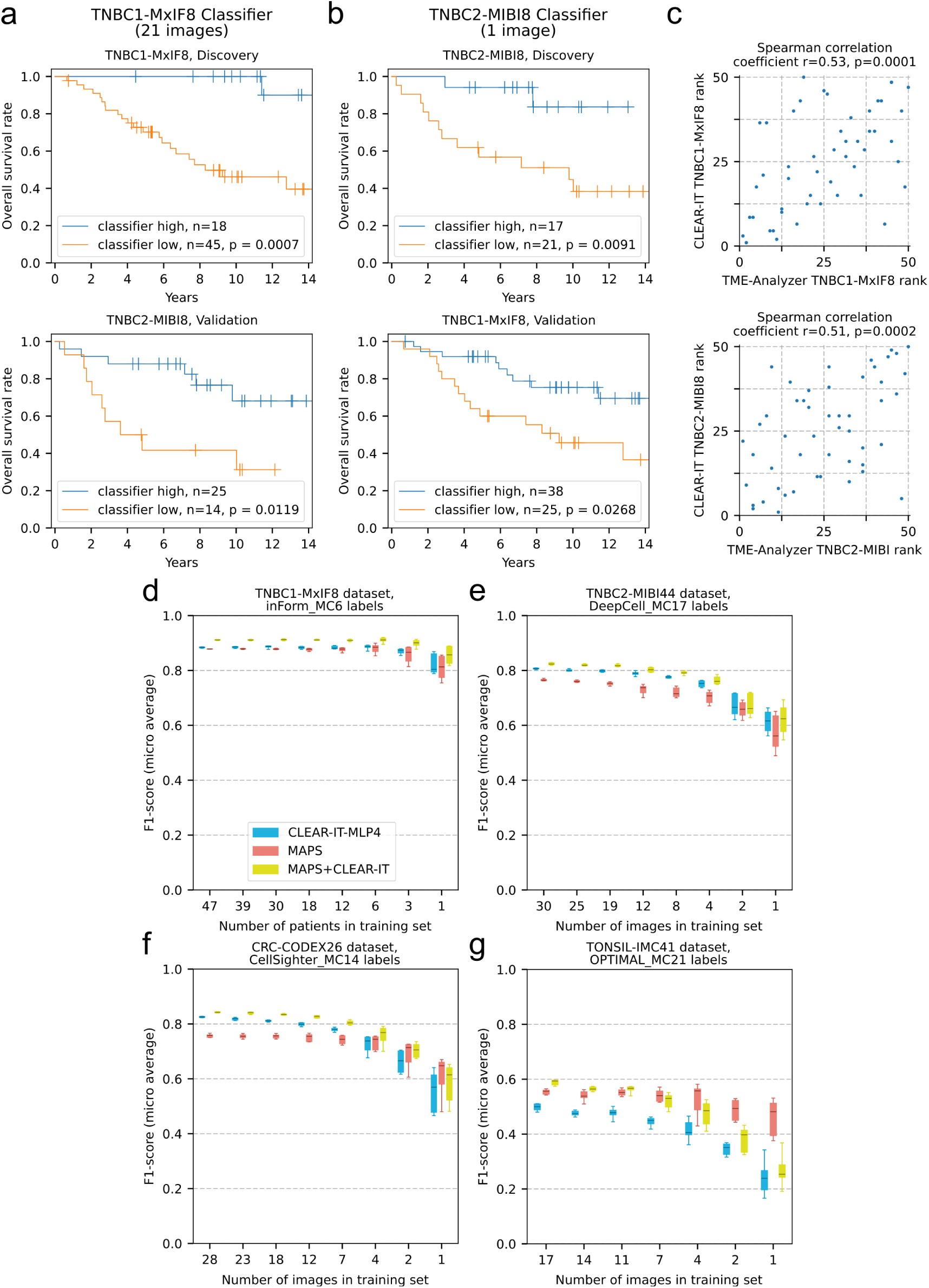
CLEAR-IT clinical and benchmark testing. **a,b)** Kaplan-Meier curves for two-group-survivals of patients in the TNBC1-MxIF8 and TNBC2-MIBI datasets based on a survival classifier built using CLEAR-IT-SLP cell classification with minimal label with training performed on discovery cohort; a: TNBC1-MxIF8 discovery (top) and TNBC2-MIBI8 validation (bottom) cohorts; b: TNBC2-MIBI8 discovery (top) and TNBC1-MxIF8 validation (bottom) cohorts. **c)** Correlation of parameter ranking based on their prognostic values in TNBC1-MxIF8 (top) and TNBC2-MIBI8 (bottom) datasets according to TME-Analyzer and CLEAR-IT analysis. **d-g)** Box-and-whisker plots showing F1 scores of CLEAR-IT-MLP4, MAPS and their combinations trained with reduced amounts of labeled data for the d) TNBC1-MxIF8, e) TNBC2-MIBI44, f) CRC-CODEX26, and g) TONSIL-IMC41 datasets with 10 boot-strappings. p-values were obtained according to log-rank test (a, b), and Spearman correlation (c).

We next benchmarked our method with MAPS (Machine learning for Analysis of Proteomics in Spatial biology) [8], where for fair comparison instead of an SLP a 4-layer multi-layer perceptron was trained with CLEAR-IT encodings (CLEAR-IT-MLP4). We also made use of two additional published datasets[12, 13], bringing the datasets tested to TNBC1-MxIF8 [6], TNBC2-MIBI44 [4], CRC-CODEX26 [12] and TONSIL-IMC41 [13]. Here, TONSIL-IMC41 differed from the other datasets, also in CLEAR-IT-MLP4 cross-performance (**Fig. E4**), as the tissue was non-cancerous and the pixel size was double (**Supplementary Information**). CLEAR-IT-MLP4 vs MAPS performance was dataset dependent: in TNBC1-MxIF8 comparable performance between two methods, in TNBC2-MIBI44 higher performance of CLEAR-IT-MLP4, in TONSIL-IMC41 higher performance of MAPS and in CRC-CODEX26 variable performance between two methods, were observed (**Fig. 2d-g**). Importantly, combination of the two methods, i.e., supervised training on the concatenation of CLEAR-IT encodings and cell expressions, consistently resulted in high performance across datasets. Together, these findings demonstrate CLEAR-IT, as a standalone platform, can be used to phenotype cells to obtain clinically relevant parameters and its combination with existing methods improves performance.

In conclusion, CLEAR-IT for the first time demonstrates that contrastive learning-based encoders can be utilized for the task of cell classification in multiplex images with very sparse labels and cell locations, irrespective of imaging and analysis platforms. CLEAR-IT was based on a relatively simple architecture that enabled the exploration of the hyperparameter space to improve its performance, and cross-dataset performance demonstrating transferability across similar datasets. Despite this, it was able to extract clinically relevant information, its classification performance was comparable to the state-of-the-art and its combination with existing model provided further performance gain. This additionally demonstrates that CLEAR-IT encoders extract information that expands beyond cell expressions. Further exploration of the CLEAR-IT training parameter hyperspace and deeper convolutional neural networks in its architecture, as well as further fine-tuning of the encoder during supervised training [10] would enhance performance at the cost of pre-training times. Irrespectively, CLEAR-IT encoders demonstrate high performance in multiplexed images analysis with highly sparse labels without segmentation masks, significantly reducing hands-on time, and can be incorporated into image analysis software QuPath [14], to obtain user-in-the-loop interface (**Fig. E5**), providing the field of digital pathology and cancer research with a robust algorithm.

## Methods

### Data pre-processing

#### Image intensity scaling

We leave all images in the same format as they are published (either 8-bit or 16-bit integer). Before feeding images to a neural network, we divide them by a constant to scale the intensity range to [0, 1]: 255 for TNBC1-MxIF8, 265 for TNBC2-MIBI44 and TNBC2-MIBI8, and 65535 for CRC-CODEX26 and TONSIL-IMC41.

#### Train/test splits

For every experiment, an identical split of data into a train and test set was used. For the considered datasets, the train/test splits are: TNBC1-MxIF8 47/15 patients (761/249 images), TNBC2-MIBI44 and TNBC2-MIBI8 30/11 patients (30/11 images), CRC-CODEX26 28/7 patients (28/7 images), TONSIL-IMC41 5/2 patients (17/7 images). For the supervised training of classifiers, the train set was further split by 80/20 into a train and validation set. To ensure that classifiers are trained and tested on a comparable amount of data for the cross-testing of TNBC1-MxIF8 and TNBC2-MIBI8 (**Fig. 1e**), the train/test split was set to 30/11 patients for both datasets. Furthermore, for TNBC1-MxIF8, only 3 images per patient (captured at the tumor center) were included. This balanced the amount of training and testing data between the TNBC1-MxIF8 images (1340 × 1008 pixels) and the TNBC2-MIBI8 images (2048 × 2048 pixels) as much as possible to approximately 4 megapixels of image data per patient.

### CLEAR-IT training pipeline

The CLEAR-IT training pipeline is summarized in **Fig. S1**. The inference pipeline (**Fig. S1a**) takes a multiplex image patch centered around a cell as input and outputs a vector containing probabilities of the cell belonging to the class labels considered. To achieve this, the multiplex image patch is first split into its individual channel components, which results in one grayscale single-channel image patch for each image channel. The grayscale image patches are then fed to the pre-trained encoder network (specifically, a ResNet encoder [15]) which outputs a feature vector. The individual channel feature vectors are concatenated and fed into a classification network, which predicts the cell’s class label.

#### Unsupervised encoder pre-training

Pre-training of the encoder network follows the SimCLR algorithm [9] and is summarized in **Fig. S1b**. The pre-training dataset consists of grayscale single-channel image patches centered around cells from individual color channels of the multiplex images. A minibatch is constructed by sampling a number of different image patches of random cell locations and random image channels. Every image patch in the minibatch is augmented in two random and distinct ways to create an augmented image pair, which are then encoded by a ResNet-18 [15] encoder. The encodings are non-linearly projected by the projection head, which is a multi-layer perceptron (MLP) with 3 hidden layers. The loss function acts on the non-linear projections of the encodings to maximize the similarity between the outputs originating from an augmented image pair (positive pair), while minimizing the similarity across projections from augmented images that are not from the same image pair (negative pairs). After pre-training, the MLP is removed, and the ResNet encoder can be used to provide feature embeddings as input for training downstream networks.

The input size to the ResNet-18 encoder is 64 × 64 × 1 pixels (32 × 32 × 1 for the TONSIL-IMC41 dataset) and the output size is either 512 or 32, for a linear classifier or an MLP classifier, respectively, which is used to perform classification. The MLP projection head has an output layer of size 128 and 3 hidden layers, each of size 512 with batch normalization and ReLU (rectified linear unit) activations. Optimization is performed via the LARS (layer-wise adaptive rate scaling) [16] method with a learning rate of 0.3*N/*256 (*N* being the pre-training batch size), and the NT-Xent (normalized-temperature cross-entropy) loss function

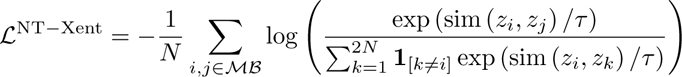

where *i, j ∈ ℳℬ* represents all augmented image pairs in the minibatch (i.e. positive pairs), *z_i_* is the non-linear feature projection of the *i* th image in the minibatch, sim (*a, b*) is the cosine similarity between vectors *a* and *b*,**1**_[_*_k≠i_*_]_ is an indicator function that is equal to 1 iff *k ≠ i* and 0 otherwise, and *τ* is the temperature parameter that is used to scale the relative importance of positive and negative pairs.

During the optimization of pre-training augmentations, we apply combinations of augmentations in the following order:

1. **Gaussian blur**: Apply Gaussian blur with intensity *x*, where *x* determines the kernel size as 2*x* + 1 and the standard deviation randomly drawn from *σ* ∈ [0*, x*].
2. **Brightness/Contrast**: Adjust brightness and/or contrast of the image by *x*, which determines the intensity that is randomly drawn from *β* ∈ [*−x, x*].
3. **Translation**: Move the center of the crop in a random radial direction. The translation distance is drawn from *ρ* ∈ [0*, x*].
4. **Zoom**: Zoom in or out of the image. The zoom factor is randomly drawn from *Z* ∈ [*x*_zmin_*, x*_zmax_] where *x* = 1 means that no zoom is applied.
5. **Rotation**: Randomly apply rotation in multiples of 90*^◦^*.
6. **Flipping**: Randomly apply horizontal and/or vertical flipping.

The optimization of the hyperparameters is illustrated in **Fig. S1d**.

#### Supervised classifier training

The supervised classifier training is the second step in the training pipeline and is summarized in **Fig. S1c**. The pre-trained encoder is used as many times in parallel as there are color channels in the input image and the computed feature encodings are concatenated and input to a classification network. For every experiment, we keep the weights of the pre-trained ResNet encoder frozen and only train the classifier to make predictions based on the generated feature embeddings.

In order to evaluate the quality of the features learned by the pre-trained encoder, we use a single-layer perceptron (SLP), in other words, a linear classifier, to make predictions about cell classes. This is referred to as “linear evaluation” [9]. The linear classifier consists of a single layer with input size 512 *× C* and output size *K*, where *C* is the number of color channels in the input image and *K* is the number of classes to predict. We train every linear classifier with a batch size of 64, a 30 % dropout layer, using the Adam optimizer with both learning rate and weight decay of 10*^−^*^4^ and minimizing the binary cross-entropy loss. For every training run, we hold out a random 20 % of the training set for validation, train for 50 epochs and choose the weights from the epoch where the classifier achieved the lowest loss on the validation set.

Since the class distribution in the used datasets is highly imbalanced, we under sampled the majority classes to ensure balanced training sets (see **Supplementary Information** for an explanation of the algorithm).

### 3-round encoder optimization

The encoder hyperparameter search for optimizing downstream classification performance is performed in 3 separate rounds. Every change in hyperparameters results in a separate ResNet encoder-linear classifier pair being trained as described previously. The optimization goal is the maximization of the classifier’s performance, which we measure with the area under the precision-recall curve (PR-AUC) and report the mean of medians for PR-AUC of each class. See **Supplementary Information** for further details.

### Cross-testing experiments

For the experiments that compare classifier performance across TNBC1-MxIF8 and TNBC2-MIBI8 and their two respective annotation sets, we use two encoders that are pre-trained on the same dataset that is used for the supervised test set, i.e. the encoders are pre-trained on TNBC1-MxIF8 dataset for the upper half of the table in **Fig. 1e** and on the TNBC2-MIBI8 dataset for the lower half of the table in **Fig. 1e**. For both encoders, the remaining hyperparameters are as follows: Pre-training batch size *N* = 256, loss function temperature *τ* = 0.05, minimum zoom factor 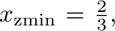 maximum zoom factor 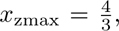 and pre-training dataset size *D* = 819200 image patches sampled from 31 patients from TNBC1-MxIF8 or TNBC2-MIBI8.

### Label-reduction by patient count

For the experiments that investigate the effects of reducing the number of patients in the supervised training set, we use the same encoders as for the cross-testing experiments, again depending on the supervised test set. For both datasets, we begin by using the maximum amount of patient data available and reduce the patient counts, resulting in classifiers trained with data from *P*_1_ ∈ {47, 39, 30, 18, 12, 6, 3, 1} patients for the TNBC1-MxIF8 dataset and *P*_2_ ∈ {30, 25, 19, 12, 8, 4, 2, 1} patients for the TNBC2-MIBI8 dataset. For every case with a reduced number of patients, ten separate classifiers are trained on image data from different combinations of randomly and independently sampled patients.

### Supervised classifier training on individual patient data

For training the supervised classifiers on image data from individual patients, we use the same encoder pre-trained on the TNBC1-MxIF8 dataset (**Fig. E2a**) and TNBC2-MIBI8 dataset (**Fig. E2b**), respectively. Since, for some patients, there are none or only very few cells of a certain phenotype, balancing the training sets based on the least frequent cells is infeasible. Instead, we set a fixed target count *T* = 500 cells per label for the amount that every class should be represented in the training sets. If the available amount is less than *T*, all available cells are included in the training set.

### Patient image quality ranking for correlation analysis

For determining the quality of images belonging to a patient, we employ the following algorithm:

1. Calculate the standard deviations of pixels across all images belonging to an individual patient for the following channels:

- CD3, CD8, CD20, CD56, CD68 channels: Standard deviation of top 0.1 % brightest pixels.
- Background channel: Standard deviation of all pixels.
2. For each patient, assign an ordinal rank from 1 to *N*, where *N* is the number of patients, based on the following criteria:

- CD3, CD8, CD20, CD56, CD68 channels: High rank for high standard deviation.
- Background channel: High rank for low standard deviation.
3. For each patient, sum the ranks assigned for each channel.
4. Divide all rank sums by the highest obtained.

The resulting rank scores range between 1*/N* for lowest image quality and 1 for highest image quality per patient.

### Relating CLEAR-IT output to clinical data

The ranking of parameters and survival classifiers were obtained as described previously [11]. In brief, 50 center parameters per patient were used, namely: densities of immune cell populations in tumor and stroma compartments (*n* = 12); areas of these compartments (*n* = 2); and distance z-scores between phenotypes in whole tissue (*n* = 36), and were ranked with a reiterative nested Monte Carlo approach. Here, the patient dataset was randomly split into train and test sets and interim classifiers were constructed in the train set. First, the representative values of parameters for long and short survival were obtained through repeating the following steps 1,000 times:

1. Random selection of 12 patients;
2. Splitting these 12 patients for each of 50 parameter into 2 groups of 6 patients according to the median value of parameters;
3. Testing statistical significance of survival differences between the 2 patient groups per parameter according to log-rank tests; and finally
4. Upon significance, recording the parameter mean values for the shorter and longer survival groups.

After 1,000 repetitions, the means of statistically significant parameters were calculated for shorter and longer survival groups, and the parameters were ranked based on the degree of separation (p-value) between the survival groups obtained in the train set. Second, the patients in the train set were split into two groups, based on most parameters for each patient being closer to the recorded means of shorter or longer survivors, where the lowest ranking parameter was reiteratively excluded until statistically significant performance was reached. Third, the surviving interim classifier (i.e., the resulting set of parameters and their recorded means) was tested in the test set. In the case when statistical significance was also observed for the test set, then the set of parameters that defined the classifier were considered to have a first hit. These three steps were repeated 5,000 times and the parameters were ranked based on the total number of hits in the interim classifiers. Subsequently, the top 10 ranking parameters were used to obtain the final survival classifier using the steps 1-4 above on the whole discovery cohort, but with a repetition of 5,000 instead of 1,000, and then applied to the discovery cohort.

### CLEAR-IT feature benchmark against MAPS algorithm

In order to benchmark the quality of feature encodings produced by a pre-trained CLEAR-IT encoder, we utilize the MAPS [8] architecture for classification. Briefly, MAPS consists of a MLP with 4 hidden layers that performs cell classification based on cell expressions and cell size, resulting in an input size of *C* +1, where *C* is the number of channels in the image, and an output size of *K*, where *K* is the number of class labels. In order to perform classification of CLEAR-IT features, we pre-train encoders to output feature vectors of size 32 and adapt the MAPS input size to be 32 *× C*. Lastly, we investigate performance of concatenating both inputs, resulting in an input size of *C ×* 33 + 1. When training MAPS on cell expressions, we use the following training parameters: batch size 128, 25 % dropout, Adam optimizer with learning rate of 10*^−^*^4^ and minimizing the binary cross-entropy loss. When training MAPS on CLEAR-IT features and their concatenation with cell expressions, we use the following training parameters: batch size 1024, 25 % dropout, Adam optimizer with learning rate of and minimizing the binary cross-entropy loss.

### Calculation of cell size and cell expressions

In order to provide an input to the MAPS architecture, we calculate the cell sizes and cell expressions from the available images. Cell sizes are determined by counting the pixels belonging to a particular cell in the segmentation masks. Cell expressions per channel are calculated by summing the pixel intensities inside segmented regions per channel and dividing this by the cell size.

### Hardware and software implementation

Code has been developed and tested on a machine with an Intel Core i9-10900X CPU, 128 GB RAM and an NVIDIA GeForce RTX 3090 GPU (24GB VRAM) within the *PyTorch Release 24.07* Docker container provided by NVIDIA Optimized Frameworks. Image augmentations are implemented using Kornia [17].

Training and inference times depend on the dataset. Encoder pre-training during the hyper-parameter optimizations took around 2 hours per network and supervised training of a linear classifier around 10 minutes. Excluding loading times, inference can be performed at a speed of about 4,500 cell image patches (of size 64 × 64 × 8 pixels) per second.

## Data Availability

- TNBC1-MxIF8 dataset [1]:

– Images and segmentation masks available upon reasonable request at https://doi.org/10.4121/126d8103-6de5-4493-a48e-5d529fef471e
- TNBC2-MIBI44 dataset [4]:

– Images and segmentation masks available to download at https://www.weizmann.ac.il/mcb/Keren/resources
- CRC-CODEX26 dataset [12]

– Images available to download at https://doi.org/10.7937/TCIA.2020.FQN0-0326
– Segmentation masks provided by authors of [7]
- TONSIL-IMC41 dataset [13]:

– Images and segmentation masks available to download at https://www.ebi.ac.uk/biostudies/bioimages/studies/S-BSST1047

All data tables, pre-trained encoder models, and supervised classifier models are available to download at https://doi.org/10.4121/ebc792ad-4767-4aef-b8ff-ae653e901e3f

## Code Availability

Code is under embargo until the peer-review process is finished and will be made available with the manuscript upon publication.

## Acknowledgments

Icons in **Fig. E5c** are licensed under CC BY 3.0: “workspace” by Universal Icons, “clock” by Salman Azzumardi.

We thank Dora Hammerl, John Martens (both from Erasmus MC, the Netherlands), Yael Amitay, Leeat Keren (both from Weizmann Institute of Science, Israel), Michael Angelo (Stanford University, USA), Christian Schürch (University of Tübingen, Germany), Garry Nolan (Stanford University, USA), Bethany Hunter, Andew Filby, George Merces (all from Newcastle University, UK), Paul Rees (Swansea University, UK), Muhammad Shaban (Brigham and Women’s Hospital, USA), Yunhao Bai (Broad Institute, USA), Huaying Qiu, Sizun Jiang (both from Beth Israel Deaconess Medical Center, USA), and Faisal Mahmood (Brigham and Women’s Hospital, USA) for making their datasets and analyses available.

We also thank members of the laboratory of tumor immunology (Erasmus MC, the Netherlands), Rolf Harkes (Netherlands Cancer Institute, the Netherlands), Thibaut Goldsborough (University of Edinburgh, UK), and Dimitri Kromm (Delft University of Technology, the Netherlands) for valuable discussions.

This work in part was funded by Erasmus MC’s Academic Center of Excellence for Tumor Immunology and Immune Therapy (ACE TI-IT) and the Wellcome Trust 223750/Z/21/Z.

For the purpose of open access, the author has applied a Creative Commons Attribution (CC BY) licence to any Author Accepted Manuscript version arising from this submission.

## Author Contributions

D.S. led the work and developed the CLEAR-IT architecture; D.S., C.S., and H.B. conceptualized the work; D.S. and H.B. curated the data, performed analyses, and prepared figures; S.K. provided technical feedback for the CLEAR-IT architecture; P.B. wrote the code for integrating CLEAR-IT into QuPath; D.S., K.P., C.S., and H.B. wrote the manuscript; R.D., C.S., and H.B. supervised the work; all authors read the manuscript and provided critical feedback.

## Open Access

This article is licensed under a Creative Commons Attribution 4.0 International License, which permits use, sharing, adaptation, distribution and reproduction in any medium or format, as long as you give appropriate credit to the original author(s) and the source, provide a link to the Creative Commons license, and indicate if changes were made. The images or other third party material in this article are included in the article’s Creative Commons license, unless indicated otherwise in a credit line to the material. If material is not included in the article’s Creative Commons license and your intended use is not permitted by statutory regulation or exceeds the permitted use, you will need to obtain permission directly from the copyright holder. To view a copy of this license, visit http://creativecommons.org/licenses/by/4.0/.

## Extended Data Figures

**Fig. E1.**
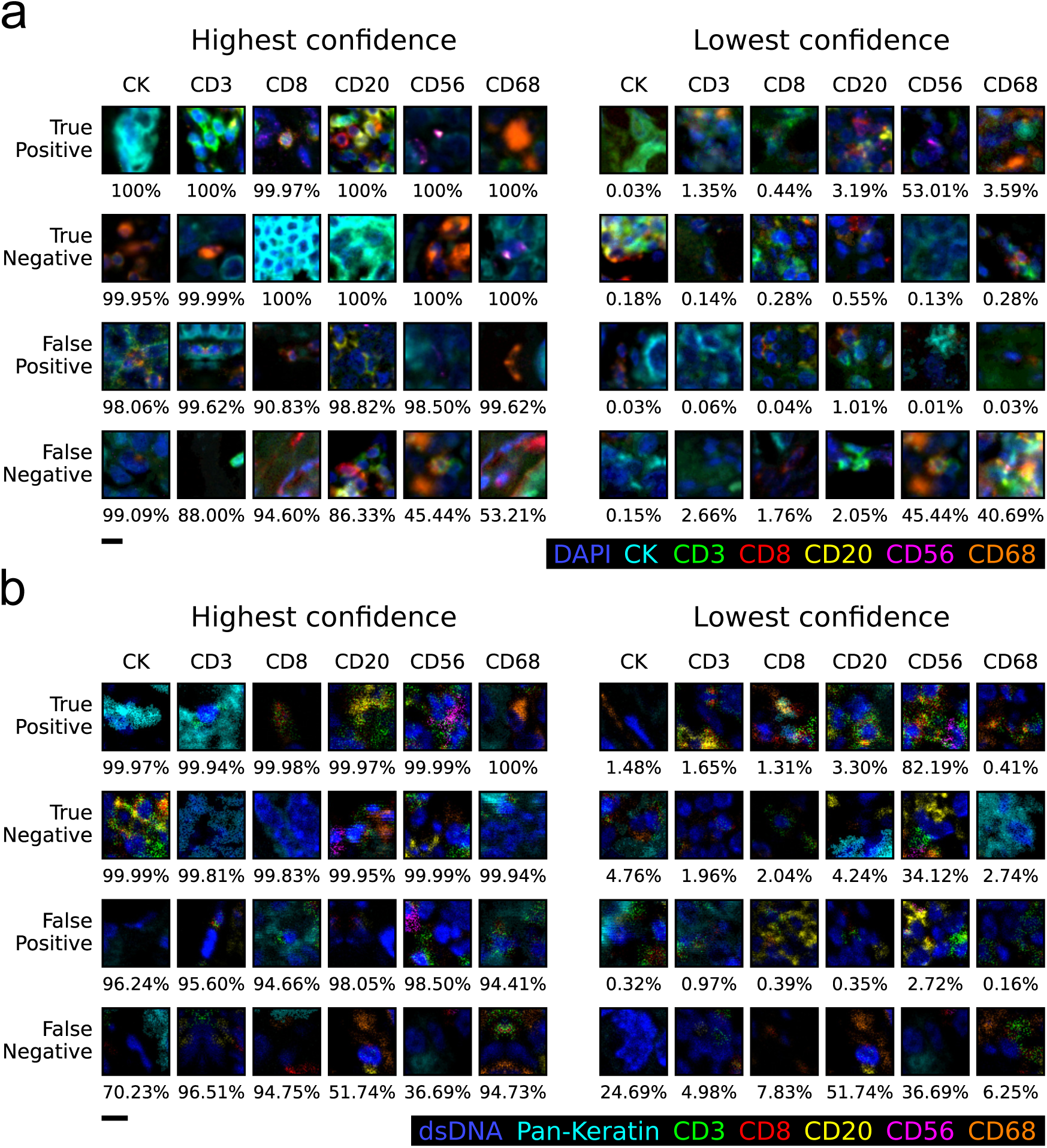
Examples of cell phenotype predictions made by CLEAR-IT. **a)** Predictions for the TNBC1-MxIF8 dataset using the best classifier obtained (3^rd^ column in **Fig. 1d**). **b)** Predictions for the TNBC2-MIBI8 dataset using the best classifier obtained (4^th^ column in **Fig. 1d**). For every type of prediction and class label, a random cell from the 100 most confident (left) and 100 least confident (right) predictions is shown. Percentage values represent the mean confidence of the 100 most and least confident predictions, respectively. For better visibility, brightness and contrast have been enhanced for the shown images. Scale bars: 10 µm.

**Fig. E2.**
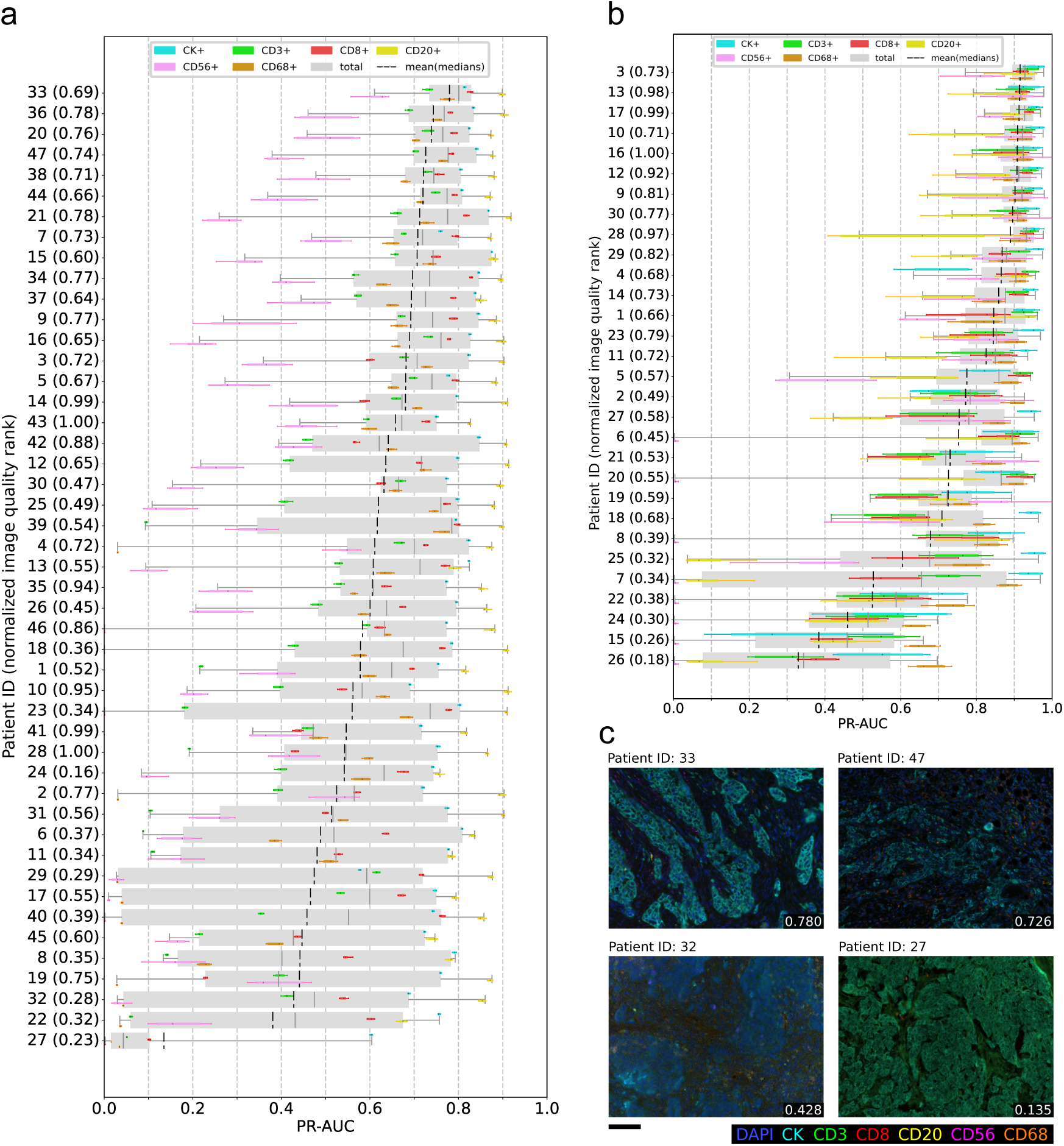
Performance of CLEAR-IT classifiers trained on data from single patients. **a,b)** Overall performance of classifiers trained on data from one patient in the TNBC1-MxIF8 (**a**) and TNBC2-MIBI8 (**b**) datasets. The area under the precision-recall-curve (PR-AUC) of the encoder/linear classifier combination, plotted per cell label and for all cells per train/test set combination. The gray boxes represent the total classification performance irrespective of class labels. The dashed black lines represent the mean of the median PR-AUC scores per class, which are represented by the colored boxes. The y-axis specifies the patient ID and the normalized image quality rank in parentheses. The boxes represent the interquartile range (25^th^ to 75^th^ percentiles) and the whiskers extend to the 5^th^ and 95^th^ percentiles of the data, excluding outliers. **c)** Example tissue images from patients with best (top row) and worst (bottom row) generalization performance in TNBC1-MxIF8 cohort, when only a single patient data is used for classifier training. Numbers in bottom right corners indicate the mean of median PR-AUC scores achieved when training classifiers on images of only that respective patient. Scale bar: 100 µm.

**Fig. E3.**
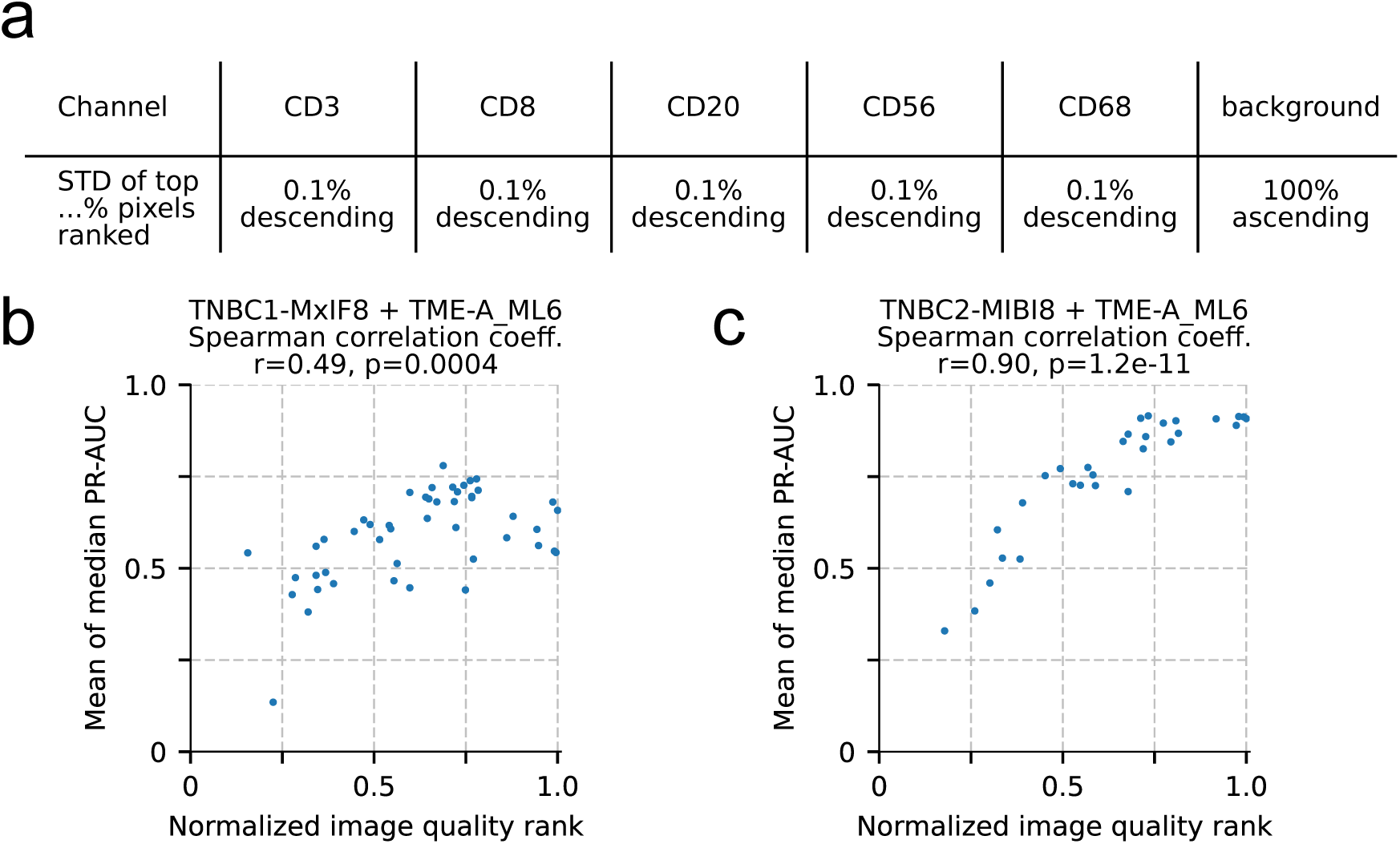
Correlation of classification performance with image quality ranking based on signal-to-noise ratio (SNR). **a)** Channel ranking parameters used for image ranking based on the pixel intensity standard deviation of highest pixel intensities. **b)** Scatter plot showing mean of median area under the precision-recall curve (PR-AUC) scores against normalized image quality rank for the TNBC1-MxIF8 dataset and TME-A ML6 annotations. Each dot represents a classifier that is trained on images from only a single patient. The corresponding normalized image quality rank is computed from images from that respective patient. **c)** Same as **b** but for the TNBC2-MIBI8 dataset and TME-A ML6 annotations.

**Fig. E4.**
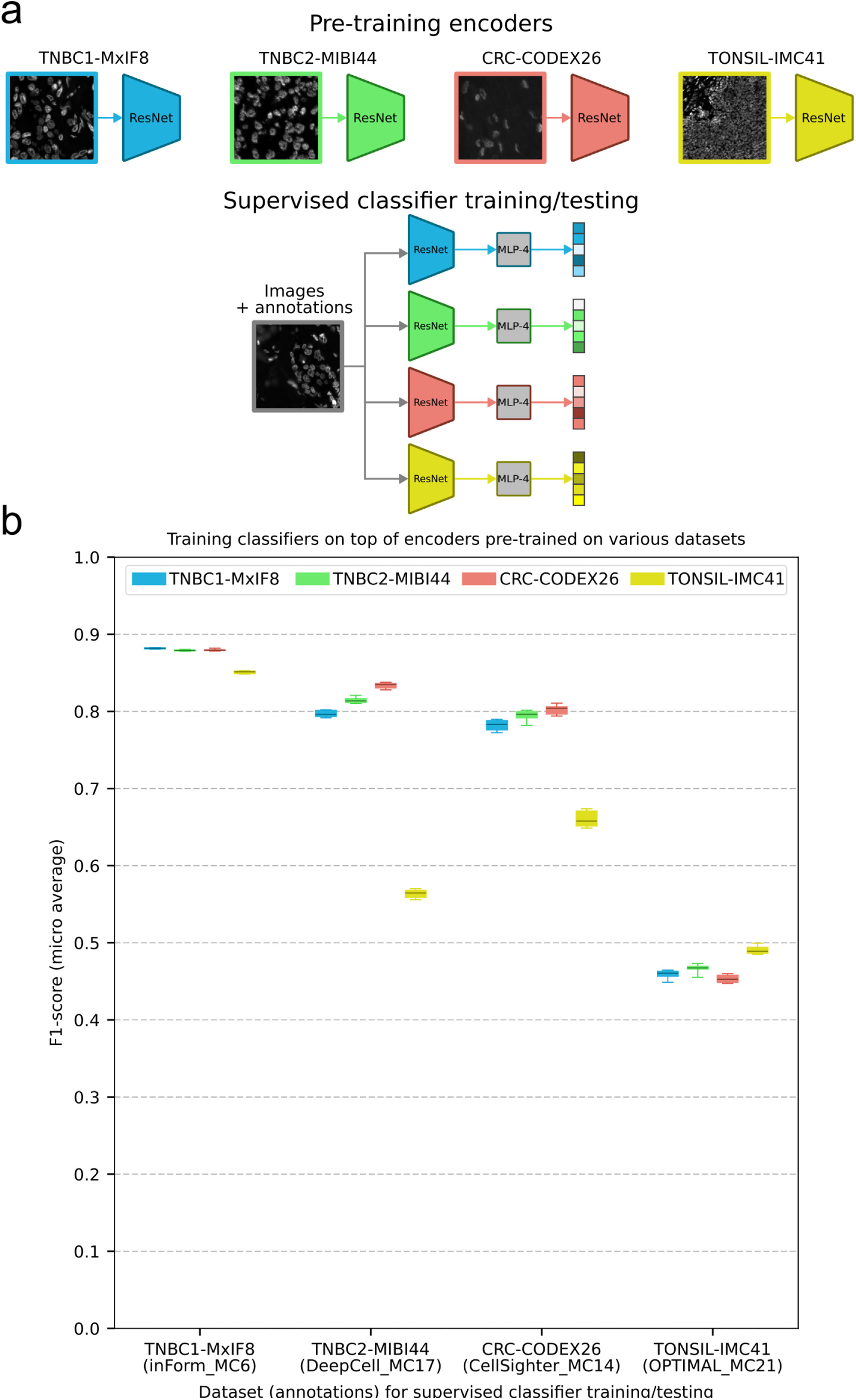
Effect of different pre-trained encoders on classification performance across datasets. **a)** For each of the four benchmark datasets (see **Fig. 2d–g**), an encoder is pre-trained (top). For the supervised training using each dataset (i.e., images and annotations), a multi-layer perceptron (MLP) with 4 layers is trained to classify the feature outputs of each of the four pre-trained encoders (bottom). This results in four classifiers per dataset for a total of 16 classifiers. **b)** Box-and-whisker plots showing micro average F1-scores of MLP classifiers as described in **a**. Each column represents one of the four benchmark datasets and the individual boxes represent which encoder was used to compute the features for the supervised training of this dataset, i.e., which dataset the encoder saw during pre-training. Every classifier was trained on the maximum amount of labeled data available for the respective dataset. The boxes represent the interquartile range (25^th^ to 75^th^ percentiles) and the whiskers extend to the 5^th^ and 95^th^ percentiles of the data, excluding outliers.

**Fig. E5.**
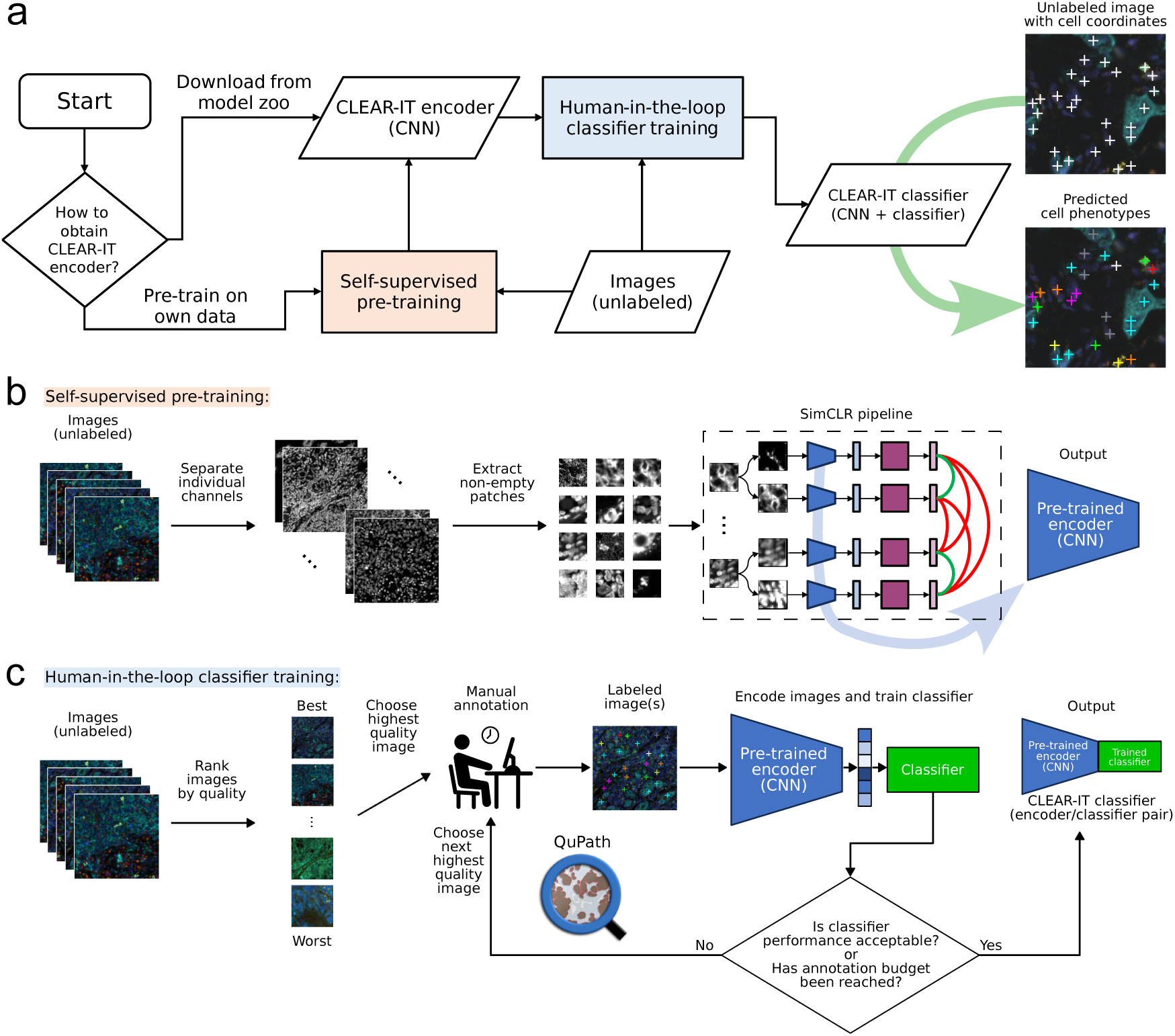
CLEAR-IT usage guideline. **a)** Flowchart showing the process from beginning with an unlabeled dataset to obtaining a CLEAR-IT classifier (encoder + classifier pair) that can perform cell phenotyping. Users can obtain a CLEAR-IT encoder either by pre-training it themselves (red box, **b**) or by downloading an already pre-trained one. The classifier on top of the encoder can be trained in a human-in-the-loop process (blue box, **c**). **b)** For the self-supervised pre-training, multiplex images are split into their individual channels and non-empty grayscale patches are extracted. The patches serve as input to the SimCLR pipeline, which produces a pretrained encoder. **c)** For the human-in-the-loop training of the cell classifier, the unlabeled images are ranked by quality. Annotation efforts are then focused on the highest quality images and iterated upon until the classification performance is acceptable or the annotation budget has been reached. The human-in-the-loop classification is facilitated by loading a pre-trained CLEAR-IT encoder into existing software, such as QuPath.

## Supplementary Information

### Datasets

#### TNBC1-MxIF8 dataset

The TNBC1-MxIF8 dataset [1] consists of 1010 multiplex immunofluorescence (MxIF) microscopy images of cancerous tissue from 62 triple-negative breast cancer (TNBC) patients. The images have a size of 1340 × 1008 pixels with pixel size of 0.5 µm, and have 8 channels: “DAPI”, “CK”, “CD3”, “CD68”, “CD8”, “CD56”, “CD20”, “background”. Overall, the images include approximately 2.3 million cells whose coordinates and cell phenotypes are specified in two separate sets of annotations, one being multi-label and the other one being multi-class. The multi-label annotation set was obtained using TME-Analyzer [11] (referred to as “TME-A ML6 labels”) and contains the following 6 classes: “CK+”, “CD3+”, “CD8+”, “CD20+”, “CD56+”, “CD68+”. The multi-class annotation set was obtained by Hammerl et al. [1] using inForm (referred to as “inForm MC7 labels”) and contains the following 7 classes: “other”, “CK”, “CD3”, “CD3 CD8”, “CD20”, “CD56”, “CD68”. Furthermore, we converted inForm MC7 to the multi-label label set inForm ML6: Here, the “CD3 CD8” class in inForm MC7 was converted to both “CD3+” and “CD8+” being positive in the multi-label equivalent.

#### TNBC2-MIBI44 dataset

The TNBC2-MIBI44 dataset [4] consists of 41 multiplex ion-beam imaging by time-of-flight (MIBI-TOF) images of cancerous tissue from 41 triple-negative breast cancer (TNBC) patients. The images have a size of 2048 × 2048 pixels with pixel size of 0.4 µm and have 44 channels: “Au”, “Background”, “Beta catenin”, “Ca”, “CD11b”, “CD11c”, “CD138”, “CD16”, “CD20”, “CD209”, “CD3”, “CD31”, “CD4”, “CD45”, “CD45RO”, “CD56”, “CD63”, “CD68”, “CD8”, “dsDNA”, “EGFR”, “Fe”, “FoxP3”, “H3K27me3”, “H3K9ac”, “HLA-DR”, “HLA Class 1”, “IDO”, “Keratin17”, “Keratin6”, “Ki67”, “Lag3”, “MPO”, “Na”, “P”, “p53”, “Pan-Keratin”, “PD-L1”, “PD1”, “phospho-S6”, “Si”, “SMA”, “Ta”, “Vimentin”. The corresponding multiclass annotation set, obtained by Keren et al. [4] (referred to as “DeepCell MC17 labels”), consists of segmentation masks for approximately 221,000 cells and contains the following 17 classes: “Unidentified”, “Tregs”, “CD4 T”, “CD8 T”, “CD3 T”, “NK”, “B”, “Neutrophils”, “Macrophages”, “DC”, “DC/Mono”, “Mono/Neu”, “Other immune”, “Endothelial”, “Mesenchymal-like”, “Tumor”, “Keratin-positive tumor”.

#### TNBC2-MIBI8 dataset

To make the images from the TNBC2-MIBI44 dataset qualitatively similar to those from the TNBC1-MxIF8 dataset, 8 out of the original 44 image channels were extracted: “dsDNA”, “Pan-Keratin”, “CD3”, “CD68”, “CD8”, “CD56”, “CD20”, “background”. Using these reduced images, a multi-label annotation set was obtained using TME-Analyzer [11] (referred to as “TME-A ML6 labels”), which contains the same 6 classes that were used for the TNBC1-MxIF8 dataset: “CK+”, “CD3+”, “CD8+”, “CD20+”, “CD56+”, “CD68+”.

To convert the DeepCell MC17 annotation set to the reduced images, the following label assignments were performed to obtain the DeepCell ML6 annotation set:

- “CD3+”: “Tregs”, “CD4 T”, “CD3 T”, “CD8 T”
- “CD8+”: “CD8 T”
- “CD56+”: “NK”
- “CD20+”: “B”
- “CD68+”: “Macrophages”
- “CK+”: “Keratin-positive tumor”

#### CRC-CODEX26 dataset

The CRC-CODEX26 dataset [12] consists of 35 co-detection by indexing (CODEX) images of cancerous tissue from 35 colorectal cancer (CRC) patients. The images have a size of 1920×1440 pixels with pixel size of 0.377 µm and have 26 channels: “CD11b”, “CD11c”, “CD15”, “CD163”, “CD20”, “CD3”, “CD31”, “CD34”, “CD38”, “CD4”, “CD45”, “CD56”, “CD57”, “CD68”, “CD8”, “Collagen”, “Cytokeratin”, “FOXP3”, “HLADR”, “MUC1”, “NAKATPASE”, “PDPN”, “SYP”, “VIM”, “SMA”, “CD45RA”. The corresponding multi-class annotation set, obtained by Amitay et al. [7] (referred to as “CellSighter MC14 labels”), consists of segmentation masks for 85,179 cells and contains the following 14 classes: “Bcell”, “CD3T”, “CD4T”, “CD8T”, “DC”, “Endothelial”, “Lymphatic”, “Macrophage”, “Neuron”, “Neutrophil”, “Plasma”, “Stroma”, “Treg”, “Tumor”.

#### TONSIL-IMC41 dataset

The TONSIL-IMC41 dataset consists of 24 imaging mass cytometry (IMC) images of human tonsil tissue from 7 individuals. The images have sizes ranging from 465 × 464 pixels to 686 × 617 pixels with pixel size of 1 µm and have 41 channels: “Sars CoV2 Spike”, “CD45RO”, “CD45RA”, “CD68”, “CD8a”, “Ki67”, “Collagen1”, “CD138”, “CD163”, “IL1R”, “CD42b”, “MPO”, “SARSCov2 Capsid”, “B7 Complement”, “CD56”, “Podoplanin”, “CD69”, “EPCAM”, “CD206”, “CD79a”, “STING”, “TMPRSS2”, “AQP5”, “CD1c”, “IFITM3”, “ACE2”, “CD57”, “p16”, “IL6R”, “cCaspase3”, “CD61”, “CD3”, “ProSPC”, “CD31”, “C30-30”, “CD4”, “HLADR”, “CD169”, “193Ir”, “CD147”, “b2M”. The images, segmentation masks, and corresponding multi-class annotation set were obtained by Hunter et al. as part of OPTIMAL (an OPTimized Imaging Mass cytometry AnaLysis framework for benchmarking segmentation and data exploration) [13]. The multi-class label set (referred to as “OPTIMAL MC21 labels”) consists of segmentation masks for 109,535 cells and contains the following 21 classes: “Epithelium”, “Memory CD8 T cells”, “M2 Macrophages”, “B cells”, “Memory CD4 T cells”, “Epithelium (proliferating)”, “Folicular B cells (proliferating)”, “Mature Macrophages”, “Folicular T cells”, “Plasma cells”, “Endothelium”, “Unclassified”, “Effector CD4 T cells”, “Epithelium and Immune cells”, “Naïve CD8 T cells”, “anti-Inflammatory Macrophages”, “CD4 T cells”, “STING+ cells”, “Apoptotic cells”, “Germinal center Macrophages”, “Standard Macrophages”.

### The design of the CLEAR-IT network architecture

In the inference pipeline (**Fig. S1a**), a multichannel image would be processed as multichannel image patches centered around the given cell locations. These image patches are split into individual channels that are fed into a pre-trained encoder, generating feature representations. For the sake of speed, in our architecture we utilized a residual neural network with 18 layers (ResNet-18) as an encoder, which is considered a small network. These representations of individual channels are concatenated into a single vector, which is then mapped to the phenotypes as probabilities through another neural network. Here, utilization of a single layer perceptron (SLP) ensured high speed and that the performance evaluated corresponded to the performance of the pre-training of the encoder on the downstream classification task, rather than the performance of a sophisticated classifier, and is often referred to as “linear evaluation” [9].

For pre-training of the ResNet-18, unsupervised contrastive learning is applied to the input images without annotations (**Fig. 1b, Fig. S1b**). Here, individual channels of multichannel image patches, for all multichannel images in the training set, are processed together as batches. Individual single-channel patches are augmented into image pairs by application of learnable non-linear transformations, encoded by ResNet, and projected to a feature representation by a non-linear multi-layer perceptron (MLP), as suggested by others [9]. This network is then trained to simultaneously maximize the similarity between the outputs of the single-channel image patch pair, and dissimilarity between outputs from a single-channel image patch and all other single-channel image patches that are not its pair. While this ResNet-18 network is smaller than others used for contrastive learning in literature [10, 18], it enabled the optimization of the hyperparameters with relatively low computational effort, and, together with the SLP, resulted in efficient inference with limited data.

After the training, the weights of the ResNet are fixed, and the SLP is trained on a train set (**Fig. S1c**). Here, the same method as the inference (**Fig. S1a**) is followed, with gradient descent to minimize the dissimilarity between the predicted probability and the true label. Upon training of the SLP, the classifier is applied to test samples, and its performance is assessed. Here, the mean of the individual class PR-AUC medians is the metric used to assess the network performance for combined evaluation of the performance of six binary classifiers on an imbalanced dataset. PR-AUC was the preferred metric as, unlike precision, recall or F1 scores, it does not require decision thresholds to be implemented and is considered more informative than the receiver operated characteristics (ROC)-AUC when evaluating binary classifiers on imbalanced datasets [8]. Additionally, usage of the mean of the individual class PR-AUC medians places equal emphasis on minority classes like CD56, instead of the overall PR-AUC median that favors majority classes like CK. This decision was made since distinct subpopulation of T cells and NK cells have been recently implicated in response to immunotherapy [19–21].

### Label conversion from multi-class to multi-label

The multi-class labels can be represented with a one-hot encoding, which is a vector where all elements are 0 except for one element which is 1. The multi-label labels can be represented with a multi-hot encoding, which is a vector where multiple elements may be 1. To convert multi-class annotation sets that explicitly define a non-positive class label (i.e. “other” for inForm MC7 and “Unidentified” for DeepCell MC17), the vector element corresponding to this class is omitted in the one-hot encoding. The resulting one-hot encoding can be directly interpreted as multi-hot encoding, where the previously explicitly defined non-positive class is now implicitly defined by all vector elements being 0.

### Undersampling algorithm for balanced multi-label training set

In order to undersample the multi-label training sets, we employ the following algorithm:

1. Class count analysis and ordering

a. Compute the frequency of each class label present in the dataset.
b. Order the classes based on their frequency from least to most frequent.
2. Inclusion of the least frequent class

a. Start with the least frequent class and add every datapoint with that class label to the training set.
b. The total number of datapoints included this way sets the target count *T*.
3. Inclusion of the more frequent classes

a. Iterate over the remaining classes from least to most frequent:

i. For each class, count the number of datapoints already included in the training set that contain the current class label. This count includes datapoints that might have been added due to their belonging to a less frequent class.
ii. Randomly sample additional datapoints from the dataset, specifically selecting datapoints that contain the current class label but are not yet in the training set, until the total count equaling to the target count *T* is reached for this class.
iii. After this sampling, all datapoints from the dataset that contain the current class label, but were not selected in this step, are permanently removed from the pool of candidates for future sampling. This step ensures to prevent the repeated selection of the same datapoints for subsequent classes.
4. Inclusion of non-positive class

a. After all positive classes have been processed, the remaining datapoints are those that do not have any positive class label assigned, e.g., “other” or “Unidentified” class.
b. From the remaining datapoints, randomly select *T* datapoints and add them to the training set.

This algorithm ensures that every class label is represented at least *T* times in the training set.

### Details on 3-round encoder optimization

During encoder optimizations, a separate PR-AUC score is computed for every class, which we report as the median obtained from applying the classifier on 10 folds of the test set. The highest mean value of those median PR-AUC scores across every class determines the best encoder whose hyperparameters are chosen to be used as default in the subsequent rounds. All pre-training is performed for 5 epochs on either the TNBC1 or TNBC2 dataset unless stated otherwise. All supervised training is performed on either the TNBC1 or TNBC2 dataset with multi-label labels and training set balancing applied. The test set is left unaltered.

#### Round 0: Base SimCLR

As a baseline, we train an encoder with the following parameters: Pre-training batch size *N* = 256, loss function temperature *τ* = 1, pre-training dataset size *D* = 409600, random rotation and flipping augmentations. Unless stated otherwise, subsequent encoders always use the same dataset size as well as the rotation and flipping augmentation.

#### Round 1: Pre-training loss function

We train encoders with pre-training batch sizes *N* ∈ {32, 64, 128, 256, 512, 1024, 2048} and loss function temperatures *τ* ∈ {0.05, 0.1, 0.5, 1, 5, 10}.

#### Round 2: Augmentations

We train encoders with translation augmentations *x*_translation_ ∈ {0, 5, 10, 15, 20, 25}, Gaussian blur augmentations *x*_blur_ ∈ {0, 2, 4, 6, 8, 10}, zoom augmentations 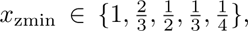 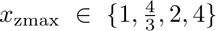 and brightness/contrast adjustment augmentations *x*_brightness_*, x*_contrast_ ∈ {0, 25, 50, 75, 100}%.

#### Round 3: Pre-training dataset size and source

We train encoders with pre-training dataset sizes *D* ∈ {409600, 819200, 2048000, 4096000} and furthermore train three encoders with dataset size *D* = 819200 and the pre-training dataset source being entirely TNBC1-MxIF8, entirely TNBC2-MIBI8 or an equal amount of data from TNBC1-MxIF8 and TNBC2-MIBI8.

## Results

We first tested the TNBC1-MxIF8 dataset[1], consisting of 1010 8-channel triple-negative breast cancer (TNBC) tissue images imaged with the Vectra multiplexed imaging system and using TME-A ML6 6 different class multi-label annotations generated with our open-source TME-Analyzer software, which we previously validated against inForm and QuPath analysis [11]. In a stepwise fashion, we optimized the encoder network by altering the pre-training loss function, augmentations used, and the training time and sample size (**Fig. S1d**), while measuring the performance of a single-layer perceptron (SLP) classifier that uses the pre-trained encoder network’s output as input. The use of a simple classifier architecture, i.e., SLP, ensures that observed performance gains can be attributed to changes made to the encoder network [9]. Here we observed the biggest benefit from the optimization of the loss function (**Fig. 1d**), where a loss temperature of *τ* = 0.05 and a pre-training batch size of 256 provided the highest performance (**Fig. S2**). We then tested different augmentations, where the value corresponds to the upper bound of a random variable used for the augmentation. Individually, the highest performance was observed for: zoom-in of 4*/*3× and zoom-out of 2*/*3×, Gaussian blur of 4 pixels, (**Fig. S3**); brightness and contrast adjustments of 0 % and 75 % (**Fig. S4**), respectively, where translation did not improve the performance (**Fig. S4**). While the highest performance gain was observed with the zoom augmentation alone, its combination with other augmentations did not further improve performance (**Fig. S5**). The augmentation parameters for the subsequent rounds were therefore set as, in addition to flipping and rotation, zoom-in of 4*/*3× and zoom-out of 2*/*3× without any other additional augmentations. Having established the parameters for the pre-training network architecture, we next tested the effect of the pre-training data on overall performance. Here, contrary to other contrastive learning tasks [9, 10], neither increasing the pre-training dataset size nor combining data from different sources improved the performance over pre-training performed on 409,600 TNBC1-MxIF8 single-channel image patches (**Fig. S6**).

We next tested our approach on the TNBC2-MIBI8 dataset [4], again using the TME-A ML6 6 class multi-label annotations. The stepwise optimization (**Fig. S1d**) again resulted in the biggest benefit from the optimization of the loss function (**Fig. 1d**), with a loss temperature of *τ* = 0.05, but with a larger pre-training batch size of 1024 (**Fig. S7**), and marginal benefit from other rounds. For augmentations tested individually; the highest performance was observed for; zoomin of 4*/*3× and zoom-out of 2*/*3×, a Gaussian blur of 0 pixels (**Fig. S8**), brightness and contrast adjustments of both 50 % (**Fig. S9**), where translation again did not improve the performance (**Fig. S9**). When combined, the highest performance was achieved for zoom-in of 4*/*3×, zoom-out of 2*/*3×, brightness adjustment of 50 %, and contrast adjustment of 50 % (**Fig. S10**), including the flipping and rotation augmentations. Hence, the only difference from the parameters used for TNBC1-MxIF8 is a larger batch size (1024 vs. 256) and the inclusion of brightness and contrast adjustments. Testing the effect of the pre-training data on overall performance, resulted in a marginal performance increase with more pre-training data, where combining data from different sources again did not further improve the performance (**Fig. S11**).

### Classifier predictions and confidence

The classifiers produce outputs using the sigmoid function, meaning that for each class, the output is a number between 0 and 1, which can be interpreted as the predicted probability of a cell belonging to that class. To obtain binary predictions, i.e., whether a prediction is positive or negative, a decision threshold is required. A prediction is considered positive if the sigmoid output exceeds this threshold, and negative otherwise. These thresholds are tuned on the held-out validation set during training to maximize the F1-score for each class.

In **Fig. E1** we present binary predictions for various classes, showing examples of true positive, true negative, false positive, and false negative predictions with the highest and lowest confidence levels, respectively. To explain how we measure this confidence, we define the confidence calculation for the model’s binary predictions. Let *y*_(*c,*pred)_ ∈ {0, 1} denote the predicted label for class *c*, where 1 represents a positive prediction and 0 a negative one. Let *σ_c_* ∈ [0, 1] be the sigmoid output (predicted probability) for class *c*, and *t_c_* ∈ (0, 1) be the decision threshold for class *c*. The prediction confidence can then be computed as follows:

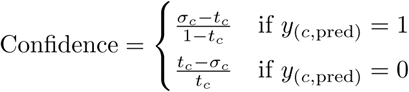

This confidence measure ranges from 0 (least confident) to 1 (most confident), providing an indication of how strongly the model’s prediction compares to the decision threshold, regardless of whether the model’s prediction is ultimately correct.

### Cross-testing of CLEAR-IT encoder across four datasets

We tested the (cross-)performance of CLEAR-IT across different datasets. For this purpose, in addition to the two TNBC datasets (TNBC1-MxIF8 [1] and TNBC2-MIBI44 [4]), we also made use of the CRC-CODEX26 [12] and TONSIL-IMC41 [13] datasets, and tested the performance of fully supervised 4-layer multi-layer perceptron (MLP) classifiers using image encodings as inputs, where the encoders were pre-trained on the images from different datasets. Here, the best and worst overall performance was achieved for TNBC1-MxIF8 and TONSIL-IMC41 dataset classifications, respectively, independent of the pre-trained encoder (**Fig. E4**). The best performance per dataset was obtained with encoders pre-trained on the same dataset as the downstream classifier, except the TNBC2-MIBI8 dataset. Best and worst generalizations were obtained for the CRC-CODEX26 pre-trained encoder and the TONSIL-IMC41 pre-trained encoder, respectively. Interestingly, encoders pre-trained on TNBC1-MxIF8, TNBC2-MIBI44, and CRC-CODEX26 all had similar performances across all datasets, together demonstrating that, while pre-training encoders on the same dataset as the downstream classifier generally improves performance, only slightly lower performance can still be achieved by encoders pre-trained on other similar datasets.

## Supplementary Figures

**Fig. S1.**
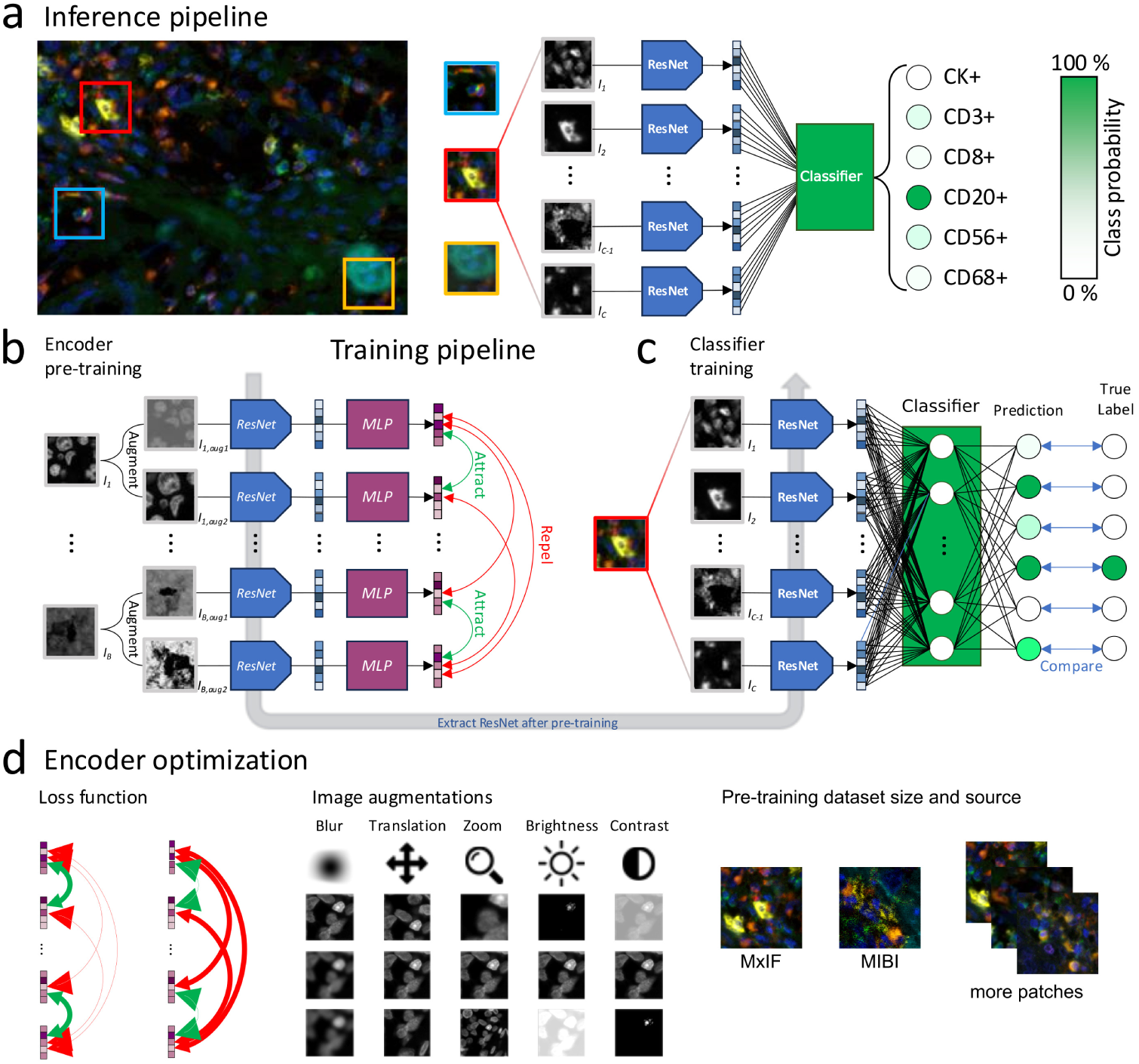
CLEAR-IT architecture and training. **a)** CLEAR-IT can be used to perform classification of individual cells in multiplex images. A patch centered around a cell of interest is split into its *C* individual image channels (from *I*_1_ to *I_C_*), each of which is processed by the same pre-trained ResNet encoder. The computed feature embeddings for each channel are concatenated and passed to a classifier, e.g. a single-layer perceptron (SLP), which predicts the class labels for the input patch. **b)** The ResNet encoder is pre-trained following the SimCLR algorithm. The inputs to the pre-training pipeline are grayscale patches which are sampled from random images, channels, and locations. Each input patch is randomly augmented in two different ways, producing an augmented image pair, which is passed to the ResNet encoder and subsequently a multi-layer perceptron (MLP). The pretraining objective is to maximize similarity between feature projections (purple squares) originating from the same image (green “Attract” arrows) and minimize similarity between feature projections that originate from a different image (red “Repel” arrows). **c)** After pre-training, the MLP is discarded and the ResNet encoder can be used to produce feature encodings for downstream tasks such as classification. By computing features for each channel of a patch in parallel and concatenating the outputs, a supervised classifier can be trained to perform classification of individual cells. **d)** The pre-training hyperparameters can be tuned to optimize the feature representations learned by the encoder. Specifically, the loss function temperature controls how strongly the maximization of similarity between feature projections is weighted, and application of image augmentations teach the encoder to become invariant to certain aberrations such that the learned feature representations contain valuable information for the intended downstream task. Increasing the size of the pre-training dataset and/or including data from different sources can improve the quality of learned features.

**Fig. S2.**
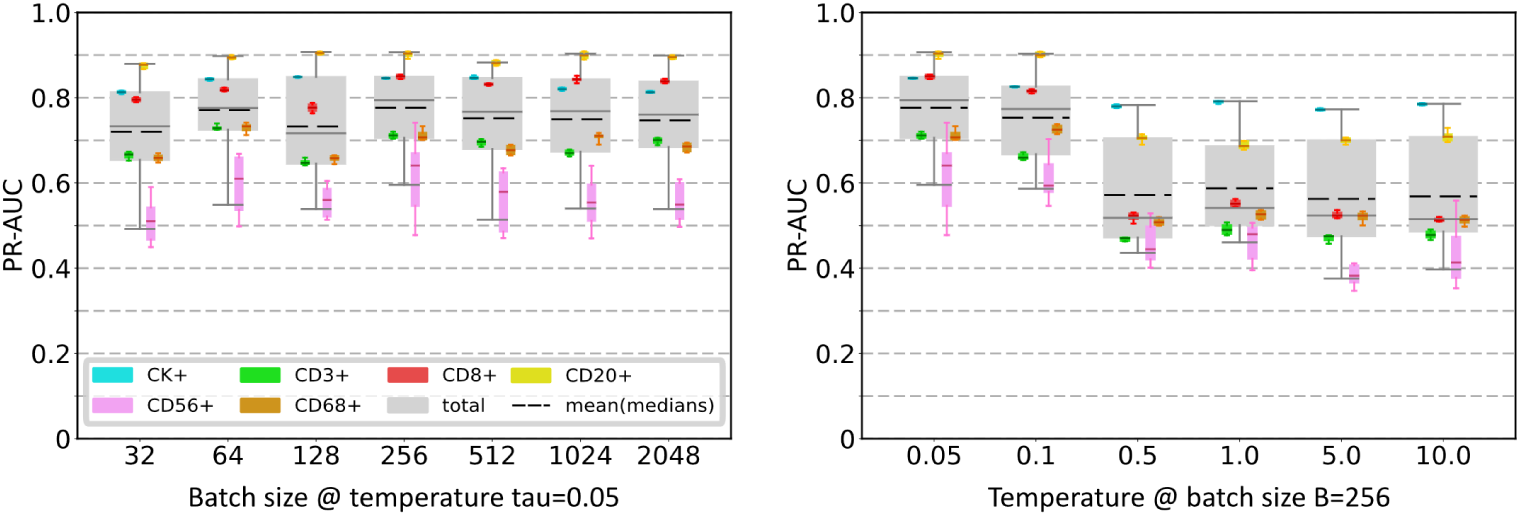
Results for round 1 of the encoder optimization for TNBC1-MxIF8 to obtain optimal temperature and batch size parameters. The area under the precision-recall-curve (PR-AUC) of the encoder/- linear classifier combination, plotted per cell type and for all cells per encoder pre-training parameter. The gray boxes represent the total classification performance irrespective of class labels. The dashed black lines represent the mean of the median PR-AUC scores per class, which are represented by the colored boxes. The encoder is pre-trained on 409,600 grayscale image patches from the TNBC1-MxIF8 dataset and the classifier is trained on the TNBC1-MxIF8 dataset with TME-Analyzer labels. The boxes represent the interquartile range (25^th^ to 75^th^ percentiles) and the whiskers extend to the 5^th^ and 95^th^ percentiles of the data, excluding outliers.

**Fig. S3.**
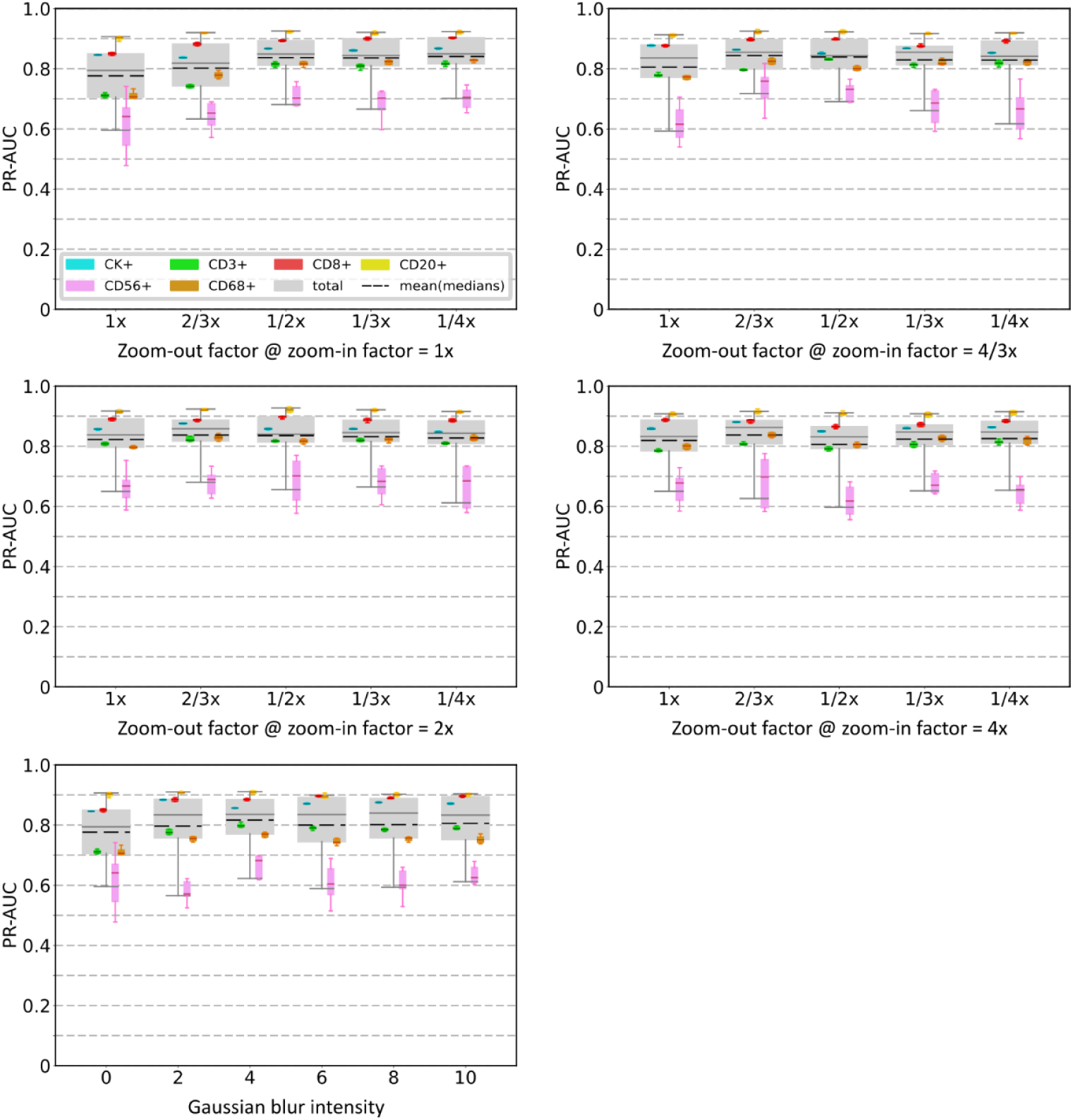
Results for round 2 (part 1 of 3) of the encoder optimization for TNBC1-MxIF8 to obtain optimal zoom and Gaussian blur parameters. The area under the precision-recall-curve (PR-AUC) of the encoder/linear classifier combination, plotted per cell type and for all cells per encoder pre-training parameter. The gray boxes represent the total classification performance irrespective of class labels. The dashed black lines represent the mean of the median PR-AUC scores per class, which are represented by the colored boxes. The encoder is pre-trained on 409,600 grayscale image patches from the TNBC1-MxIF8 dataset and the classifier is trained on the TNBC1-MxIF8 dataset with TME-Analyzer labels. The boxes represent the interquartile range (25^th^ to 75^th^ percentiles) and the whiskers extend to the 5^th^ and 95^th^ percentiles of the data, excluding outliers.

**Fig. S4.**
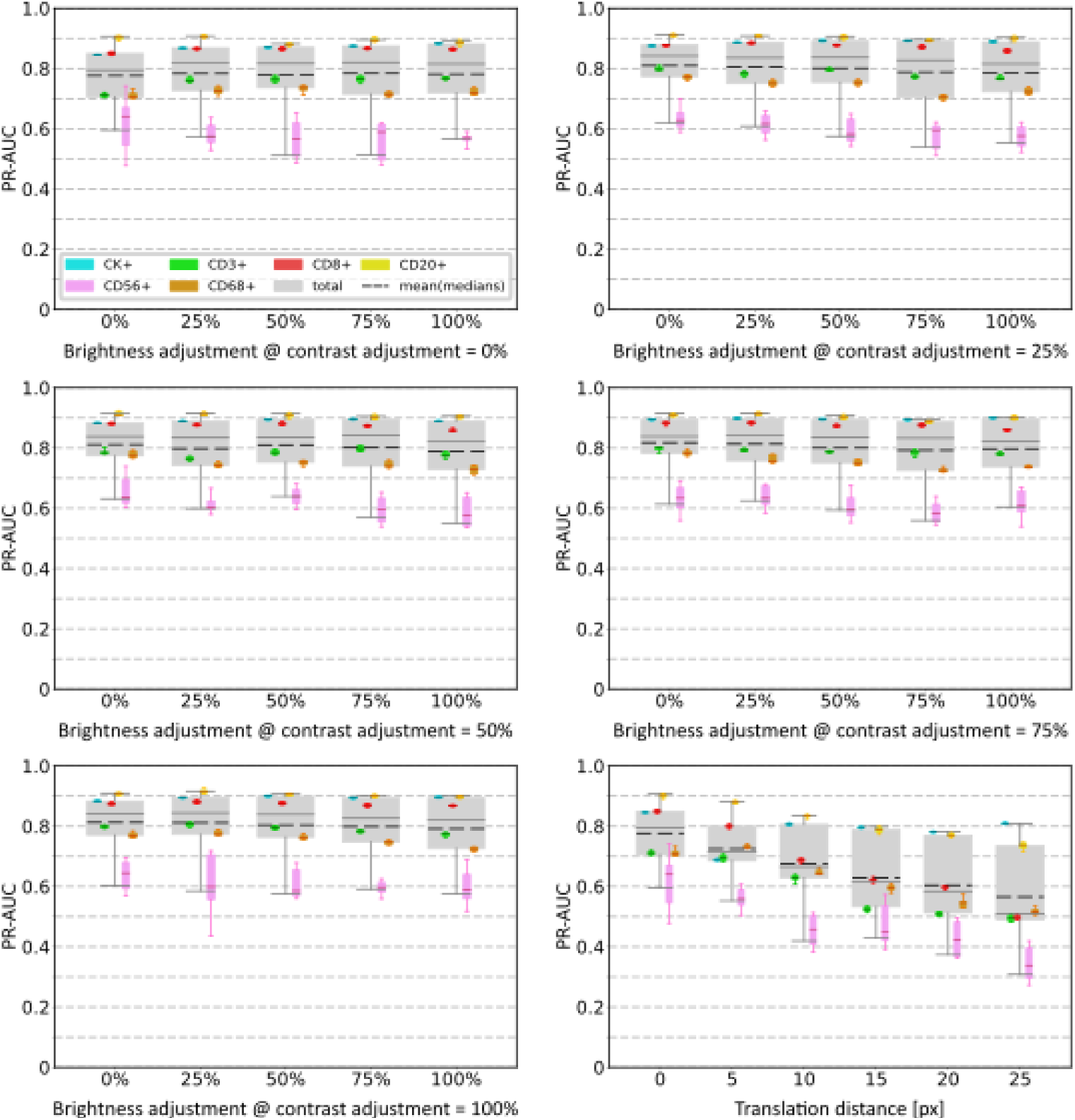
Results for round 2 (part 2 of 3) of the encoder optimization for TNBC1-MxIF8 to obtain optimal brightness/contrast and translation parameters. The area under the precision-recall-curve (PR-AUC) of the encoder/linear classifier combination, plotted per cell type and for all cells per encoder pre-training parameter. The gray boxes represent the total classification performance irrespective of class labels. The dashed black lines represent the mean of the median PR-AUC scores per class, which are represented by the colored boxes. The encoder is pre-trained on 409,600 grayscale image patches from the TNBC1-MxIF8 dataset and the classifier is trained on the TNBC1-MxIF8 dataset with TME-Analyzer labels. The boxes represent the interquartile range (25^th^ to 75^th^ percentiles) and the whiskers extend to the 5^th^ and 95^th^ percentiles of the data, excluding outliers.

**Fig. S5.**
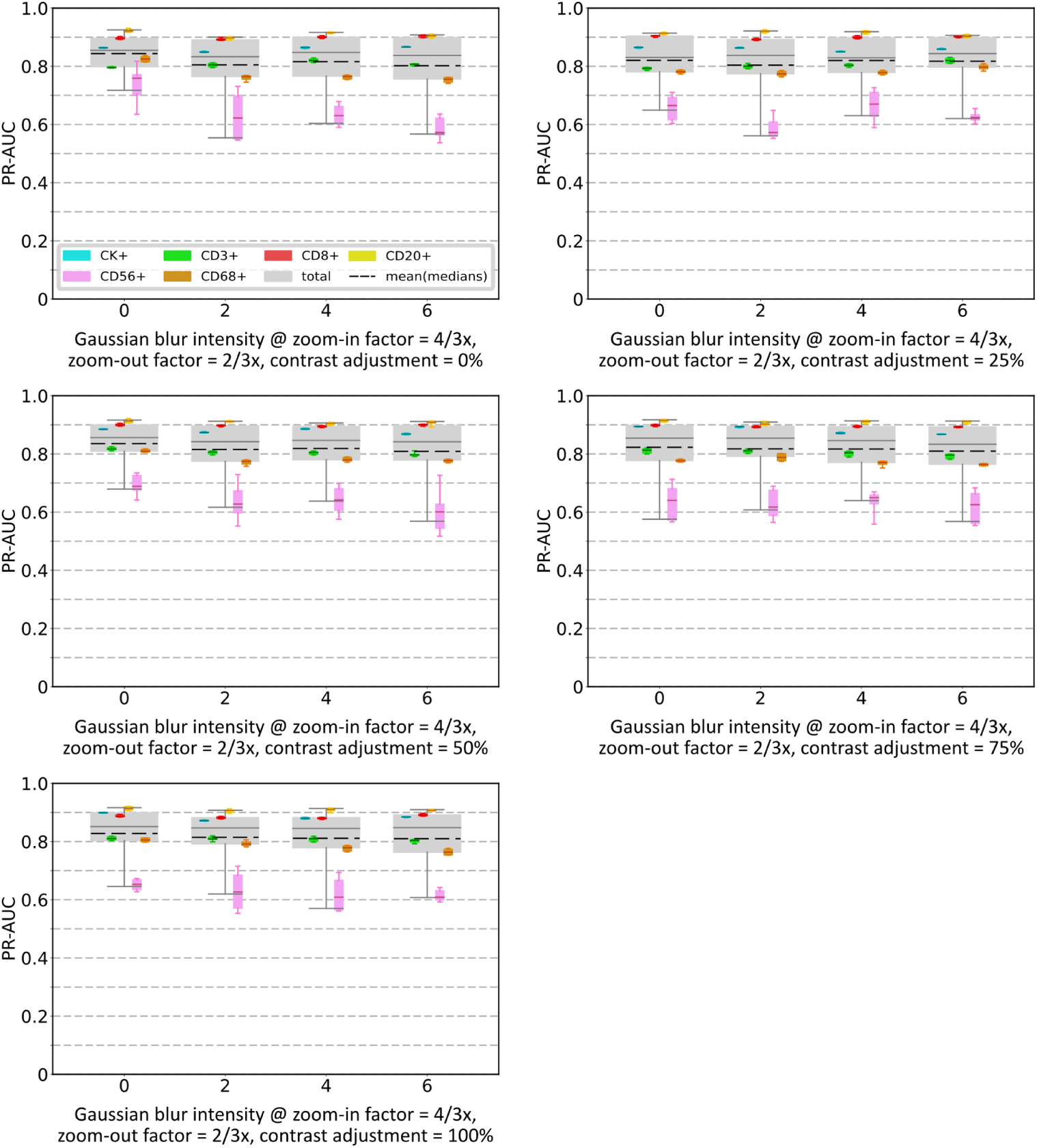
Results for round 2 (part 3 of 3) of the encoder optimization for TNBC1-MxIF8 to obtain optimal combined parameters. The area under the precision-recall-curve (PR-AUC) of the encoder/linear classifier combination, plotted per cell type and for all cells per encoder pre-training parameter. The gray boxes represent the total classification performance irrespective of class labels. The dashed black lines represent the mean of the median PR-AUC scores per class, which are represented by the colored boxes. The encoder is pre-trained on 409,600 grayscale image patches from the TNBC1-MxIF8 dataset and the classifier is trained on the TNBC1-MxIF8 dataset with TME-Analyzer labels. The boxes represent the interquartile range (25^th^ to 75^th^ percentiles) and the whiskers extend to the 5^th^ and 95^th^ percentiles of the data, excluding outliers.

**Fig. S6.**
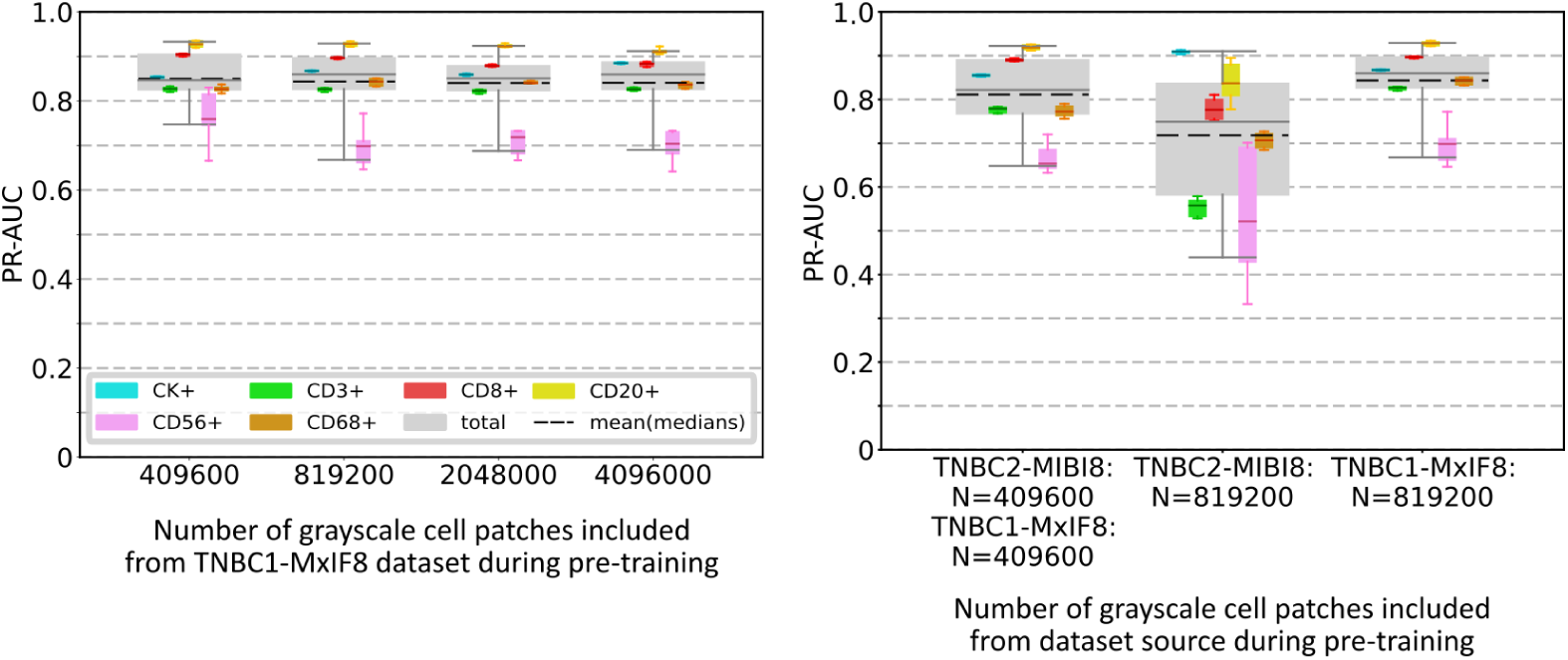
Results for round 3 of the encoder optimization for TNBC1-MxIF8 to obtain optimal pretraining data size and composition. The area under the precision-recall-curve (PR-AUC) of the encoder/linear classifier combination, plotted per cell type and for all cells per encoder pre-training parameter. The gray boxes represent the total classification performance irrespective of class labels. The dashed black lines represent the mean of the median PR-AUC scores per class, which are represented by the colored boxes. The encoder is pre-trained on grayscale image patches from the specified datasets and the classifier is trained on the TNBC1-MxIF8 dataset with TME-Analyzer labels. The boxes represent the interquartile range (25^th^ to 75^th^ percentiles) and the whiskers extend to the 5^th^ and 95^th^ percentiles of the data, excluding outliers.

**Fig. S7.**
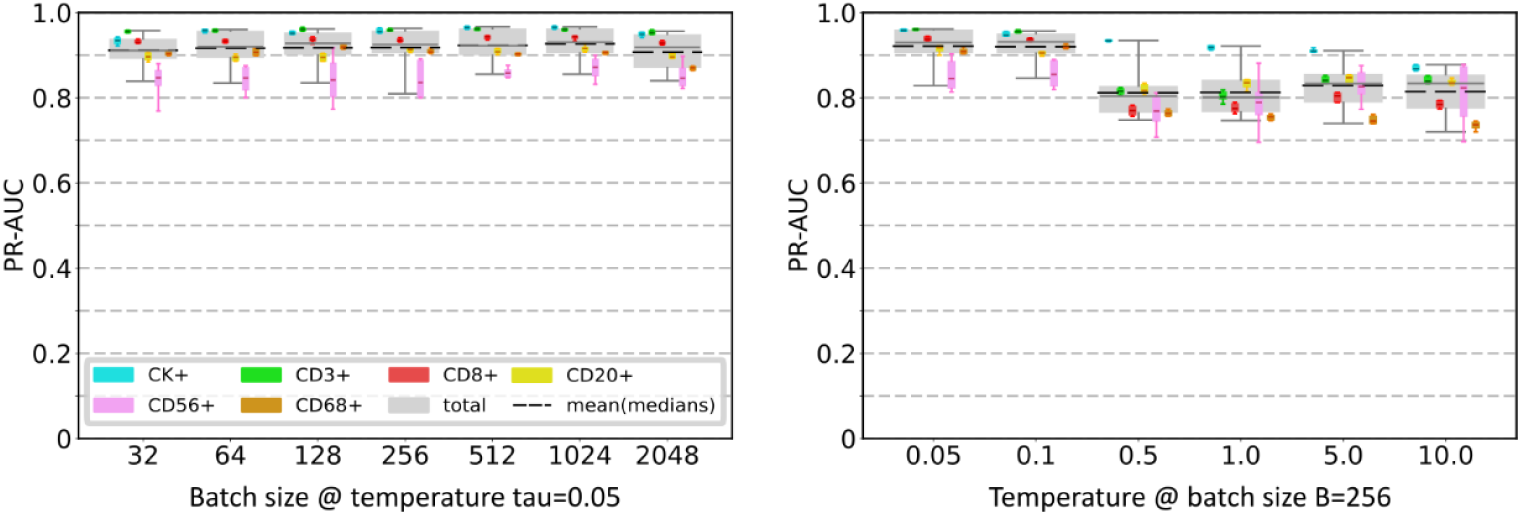
Results for round 1 of the encoder optimization for TNBC2-MIBI8 to obtain optimal temperature and batch size parameters. The area under the precision-recall-curve (PR-AUC) of the encoder/- linear classifier combination, plotted per cell type and for all cells per encoder pre-training parameter. The gray boxes represent the total classification performance irrespective of class labels. The dashed black lines represent the mean of the median PR-AUC scores per class, which are represented by the colored boxes. The encoder is pre-trained on 409,600 grayscale image patches from the TNBC2-MIBI8 dataset and the classifier is trained on the TNBC2-MIBI8 dataset with TME-Analyzer labels. The boxes represent the interquartile range (25^th^ to 75^th^ percentiles) and the whiskers extend to the 5^th^ and 95^th^ percentiles of the data, excluding outliers.

**Fig. S8.**
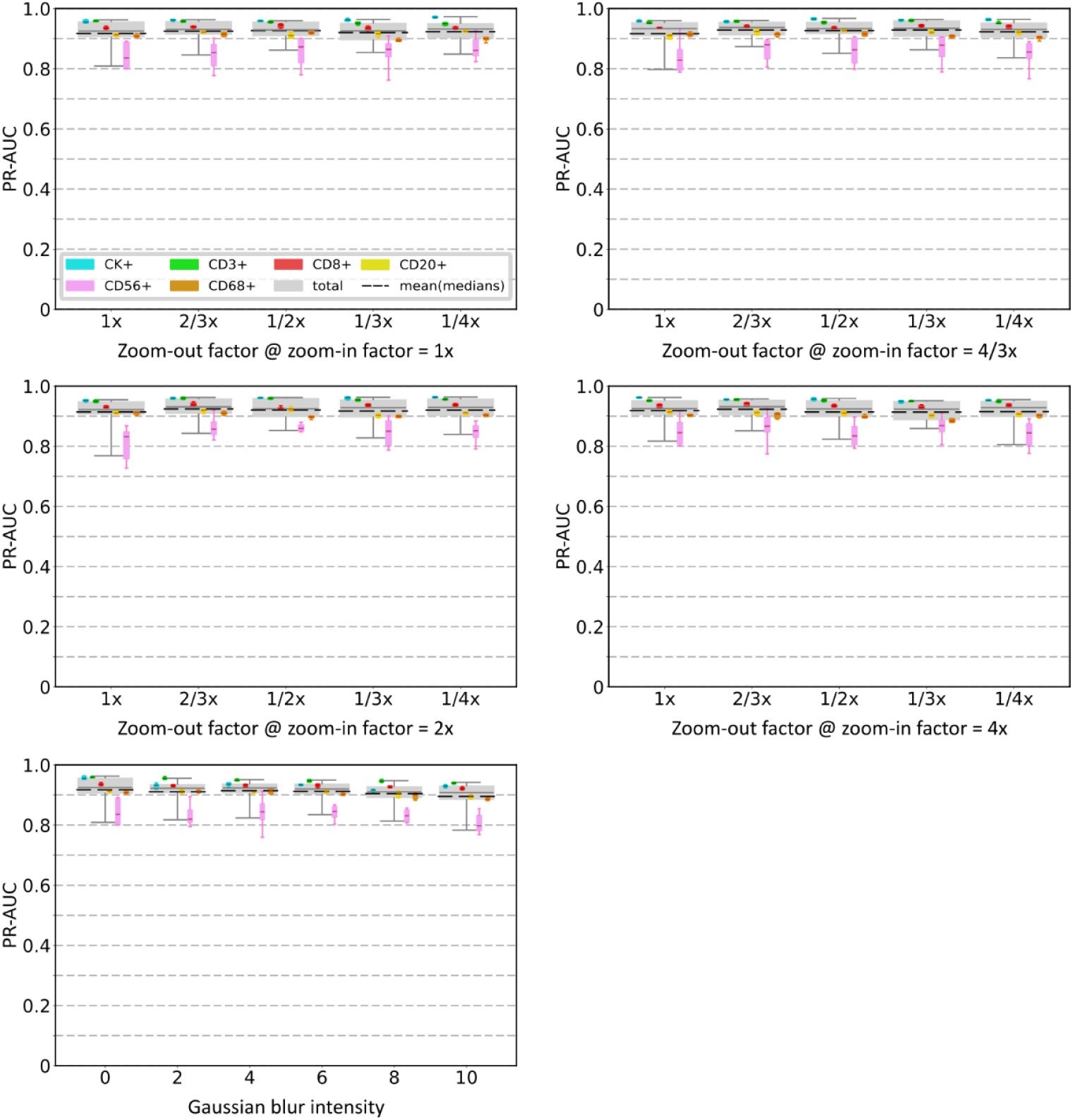
Results for round 2 (part 1 of 3) of the encoder optimization for TNBC2-MIBI8 to obtain optimal zoom and Gaussian blur parameters. The area under the precision-recall-curve (PR-AUC) of the encoder/linear classifier combination, plotted per cell type and for all cells per encoder pre-training parameter. The gray boxes represent the total classification performance irrespective of class labels. The dashed black lines represent the mean of the median PR-AUC scores per class, which are represented by the colored boxes. The encoder is pre-trained on 409,600 grayscale image patches from the TNBC2-MIBI8 dataset and the classifier is trained on the TNBC2-MIBI8 dataset with TME-Analyzer labels. The boxes represent the interquartile range (25^th^ to 75^th^ percentiles) and the whiskers extend to the 5^th^ and 95^th^ percentiles of the data, excluding outliers.

**Fig. S9.**
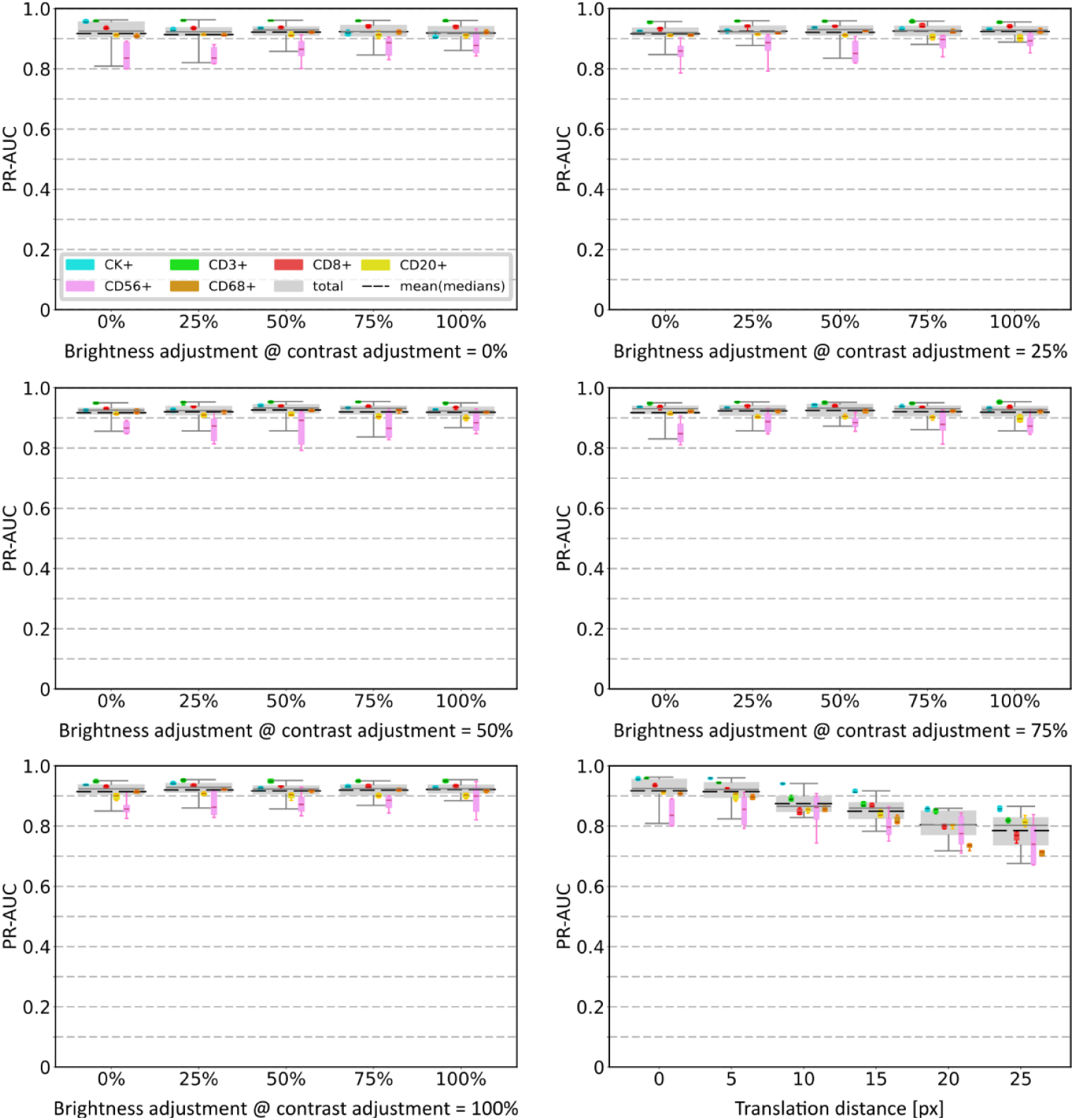
Results for round 2 (part 2 of 3) of the encoder optimization for TNBC2-MIBI8 to obtain optimal brightness/contrast and translation parameters. The area under the precision-recall-curve (PR-AUC) of the encoder/linear classifier combination, plotted per cell type and for all cells per encoder pre-training parameter. The gray boxes represent the total classification performance irrespective of class labels. The dashed black lines represent the mean of the median PR-AUC scores per class, which are represented by the colored boxes. The encoder is pre-trained on 409,600 grayscale image patches from the TNBC2-MIBI8 dataset and the classifier is trained on the TNBC2-MIBI8 dataset with TME-Analyzer labels. The boxes represent the interquartile range (25^th^ to 75^th^ percentiles) and the whiskers extend to the 5^th^ and 95^th^ percentiles of the data, excluding outliers.

**Fig. S10.**
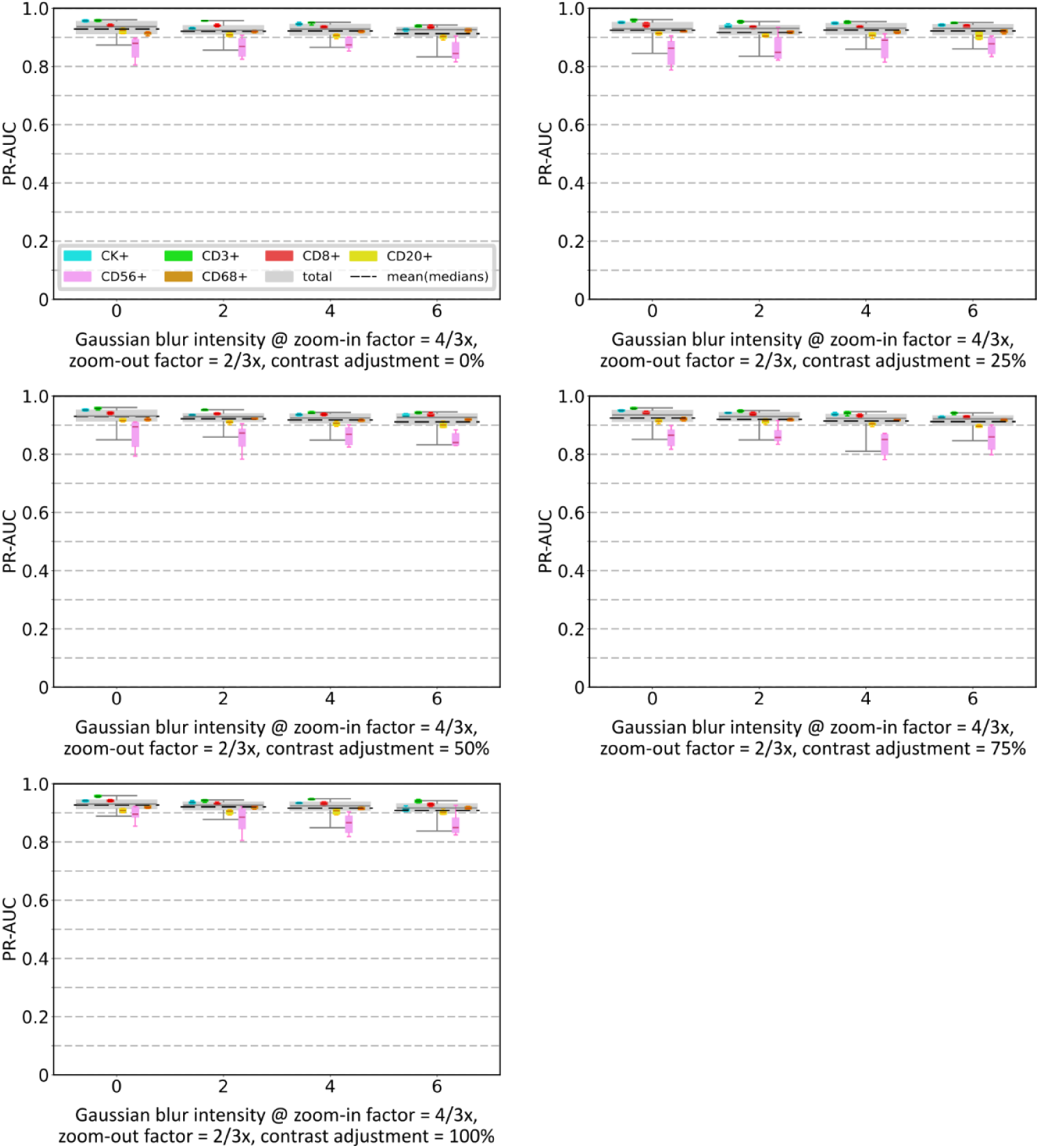
Results for round 2 (part 3 of 3) of the encoder optimization for TNBC2-MIBI8 to obtain optimal combined parameters. The area under the precision-recall-curve (PR-AUC) of the encoder/linear classifier combination, plotted per cell type and for all cells per encoder pre-training parameter. The gray boxes represent the total classification performance irrespective of class labels. The dashed black lines represent the mean of the median PR-AUC scores per class, which are represented by the colored boxes. The encoder is pre-trained on 409,600 grayscale image patches from the TNBC2-MIBI8 dataset and the classifier is trained on the TNBC2-MIBI8 dataset with TME-Analyzer labels. The boxes represent the interquartile range (25^th^ to 75^th^ percentiles) and the whiskers extend to the 5^th^ and 95^th^ percentiles of the data, excluding outliers.

**Fig. S11.**
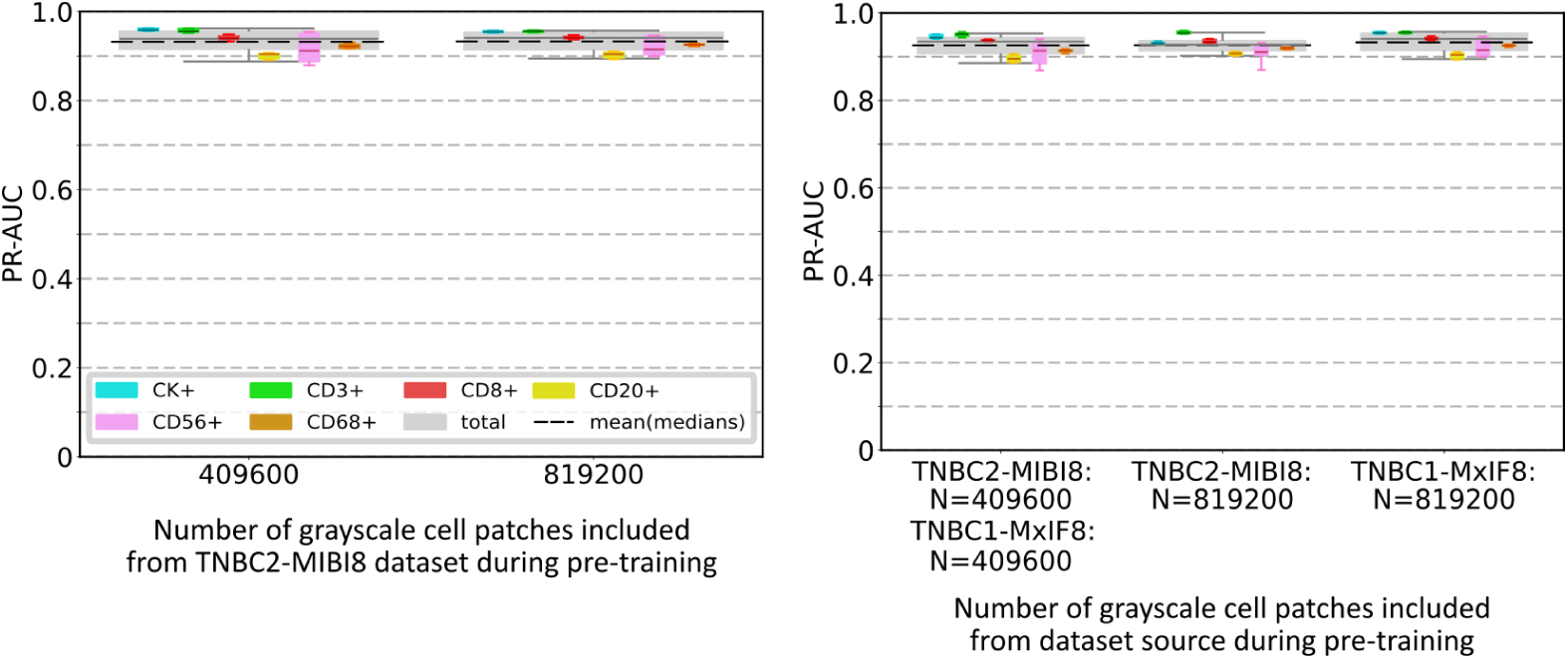
Results for round 3 of the encoder optimization for TNBC2-MIBI8 to obtain optimal pretraining data size and composition. The area under the precision-recall-curve (PR-AUC) of the encoder/linear classifier combination, plotted per cell type and for all cells per encoder pre-training parameter. The gray boxes represent the total classification performance irrespective of class labels. The dashed black lines represent the mean of the median PR-AUC scores per class, which are represented by the colored boxes. The encoder is pre-trained on grayscale image patches from the specified datasets and the classifier is trained on the TNBC2-MIBI8 dataset with TME-Analyzer labels. The boxes represent the interquartile range (25^th^ to 75^th^ percentiles) and the whiskers extend to the 5^th^ and 95^th^ percentiles of the data, excluding outliers.

**Fig. S12.**
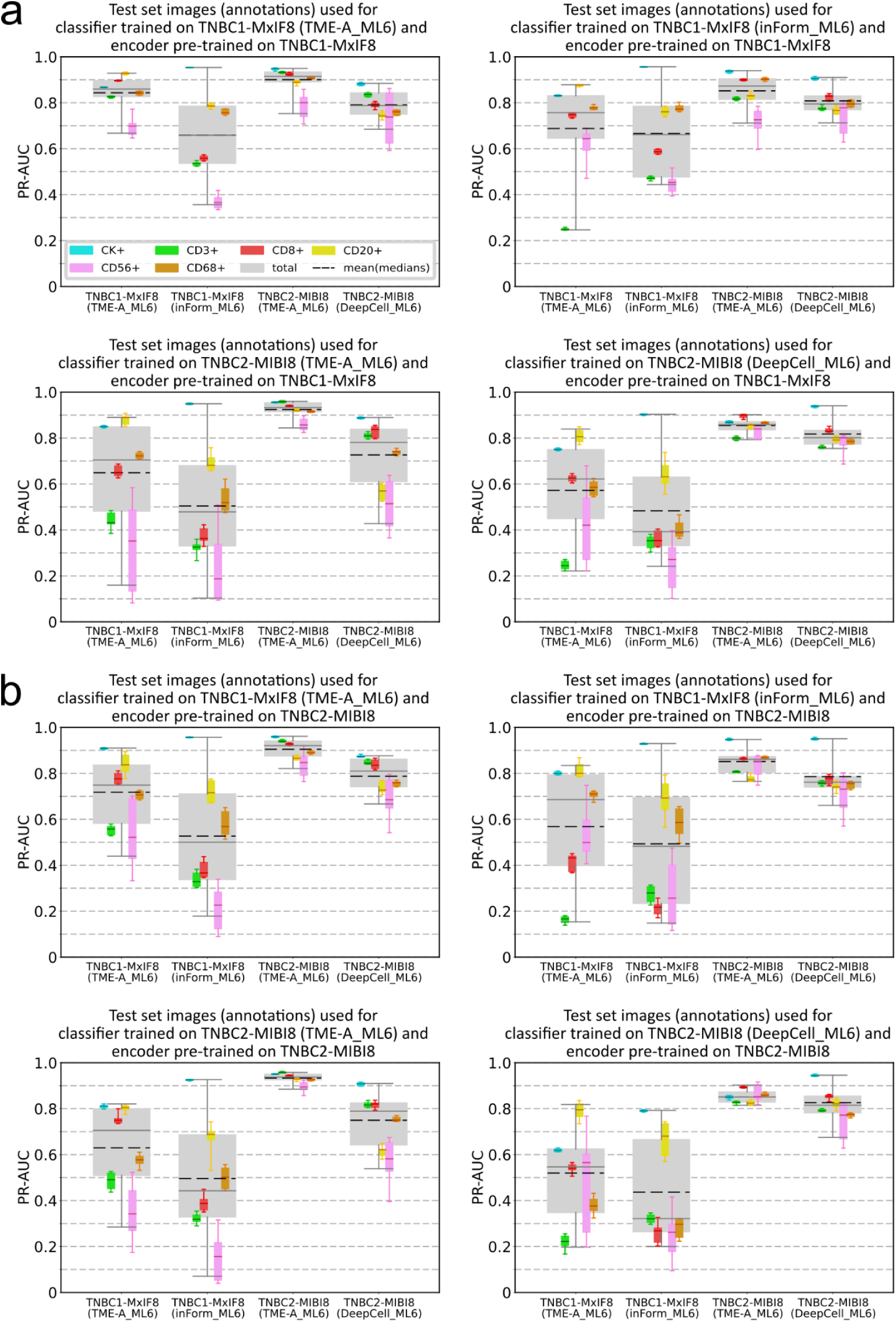
Results for cross-testing the classifiers for combinations of the two TNBC datasets. **a, b)** Classifier performance on TNBC1-MxIF8-TME-A ML6, TNBC1-MxIF8-inForm ML6, TNBC2-MIBI8-TME-A ML6, and TNBC2-MIBI8-DeepCell ML6 datasets-annotations using an encoder pre-trained on 409,600 grayscale image patches from the TNBC1-MxIF8 (**a**) or TNBC2-MIBI8 (**b**) datasets. The area under the precision-recall-curve (PR-AUC) of the encoder/linear classifier combination, plotted per cell type and for all cells per train/test set combination. The gray boxes represent the total classification performance irrespective of class labels. The dashed black lines represent the mean of the median PR-AUC scores per class, which are represented by the colored boxes. The boxes represent the interquartile range (25^th^ to 75^th^ percentiles) and the whiskers extend to the 5^th^ and 95^th^ percentiles of the data, excluding outliers.

**Fig. S13.**
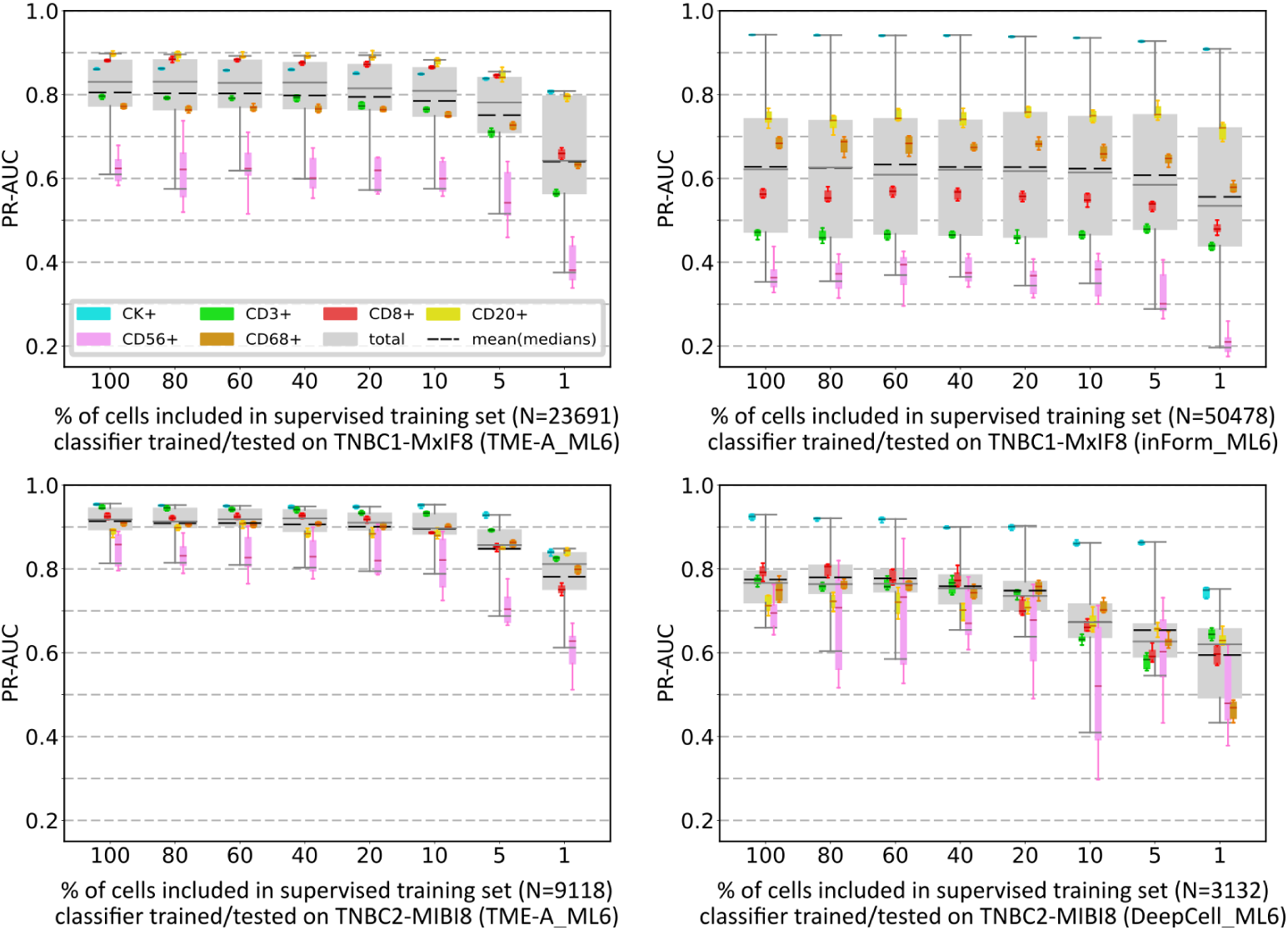
Results for label-reduction based on cell counts (irrespective of patients). The area under the precision-recall-curve (PR-AUC) of the encoder/linear classifier combination, plotted per cell label and for all cells at different percentage of labeled data used for supervised learning. Encoders are pre-trained on the same datasets as the linear classifiers. The gray boxes represent the total classification performance irrespective of class labels. The dashed black lines represent the mean of the median PR-AUC scores per class, which are represented by the colored boxes. Cell percentages are based on the maximum number of cells available in the training sets, denoted by N in parentheses. The boxes represent the interquartile range (25^th^ to 75^th^ percentiles) and the whiskers extend to the 5^th^ and 95^th^ percentiles of the data, excluding outliers.

**Fig. S14.**
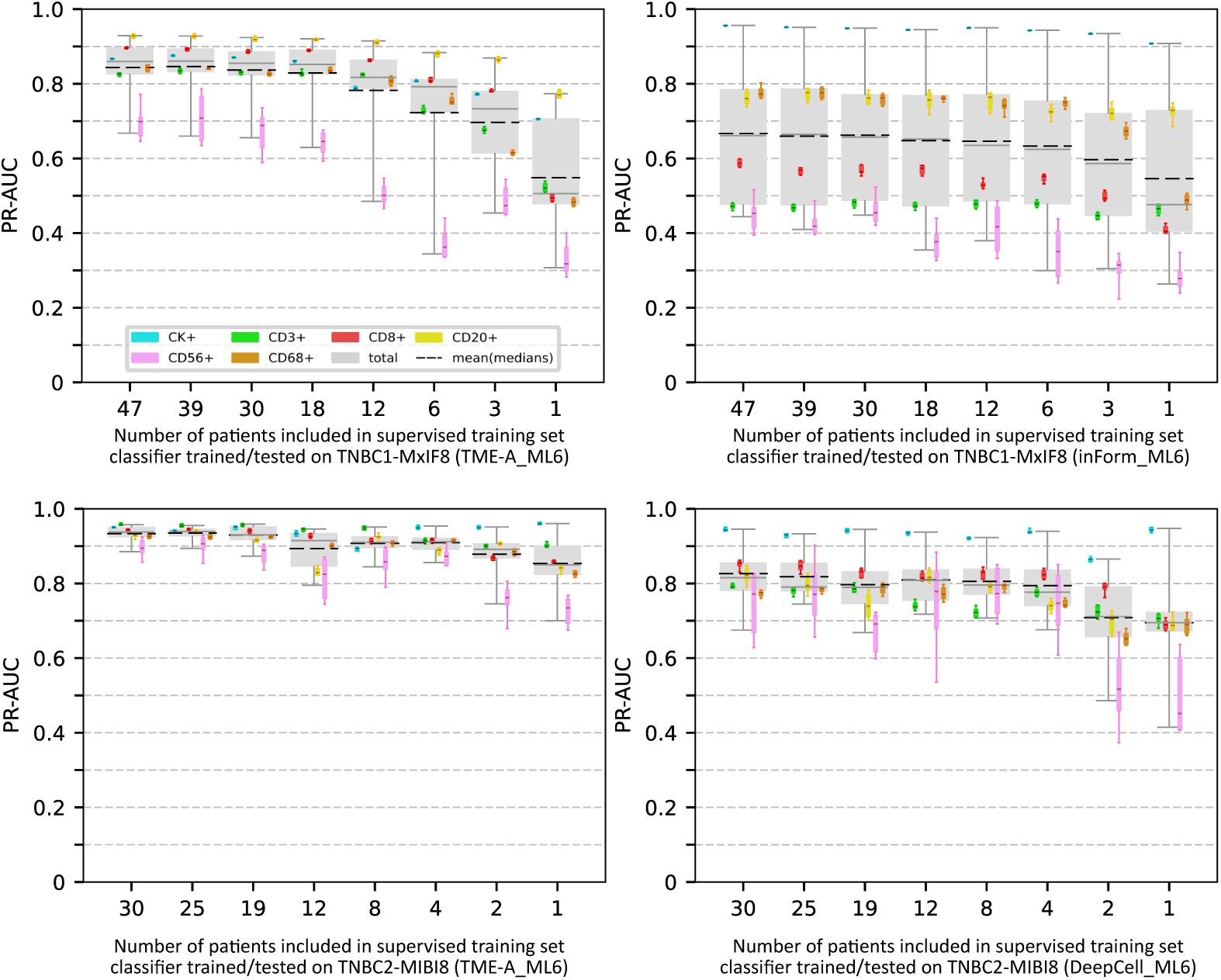
Results for label-reduction based on number of patients. The area under the precision-recall-curve (PR-AUC) of the encoder/linear classifier combination, plotted per cell label and for all cells at different percentage of labeled data used for supervised learning. Encoders are pre-trained on the same datasets as the linear classifiers. The gray boxes represent the total classification performance irrespective of class labels. The dashed black lines represent the mean of the median PR-AUC scores per class, which are represented by the colored boxes. Cell percentages are based on the maximum number of cells available in the training sets, denoted by N in parentheses. The boxes represent the interquartile range (25^th^ to 75^th^ percentiles) and the whiskers extend to the 5^th^ and 95^th^ percentiles of the data, excluding outliers.

**Fig. S15.**
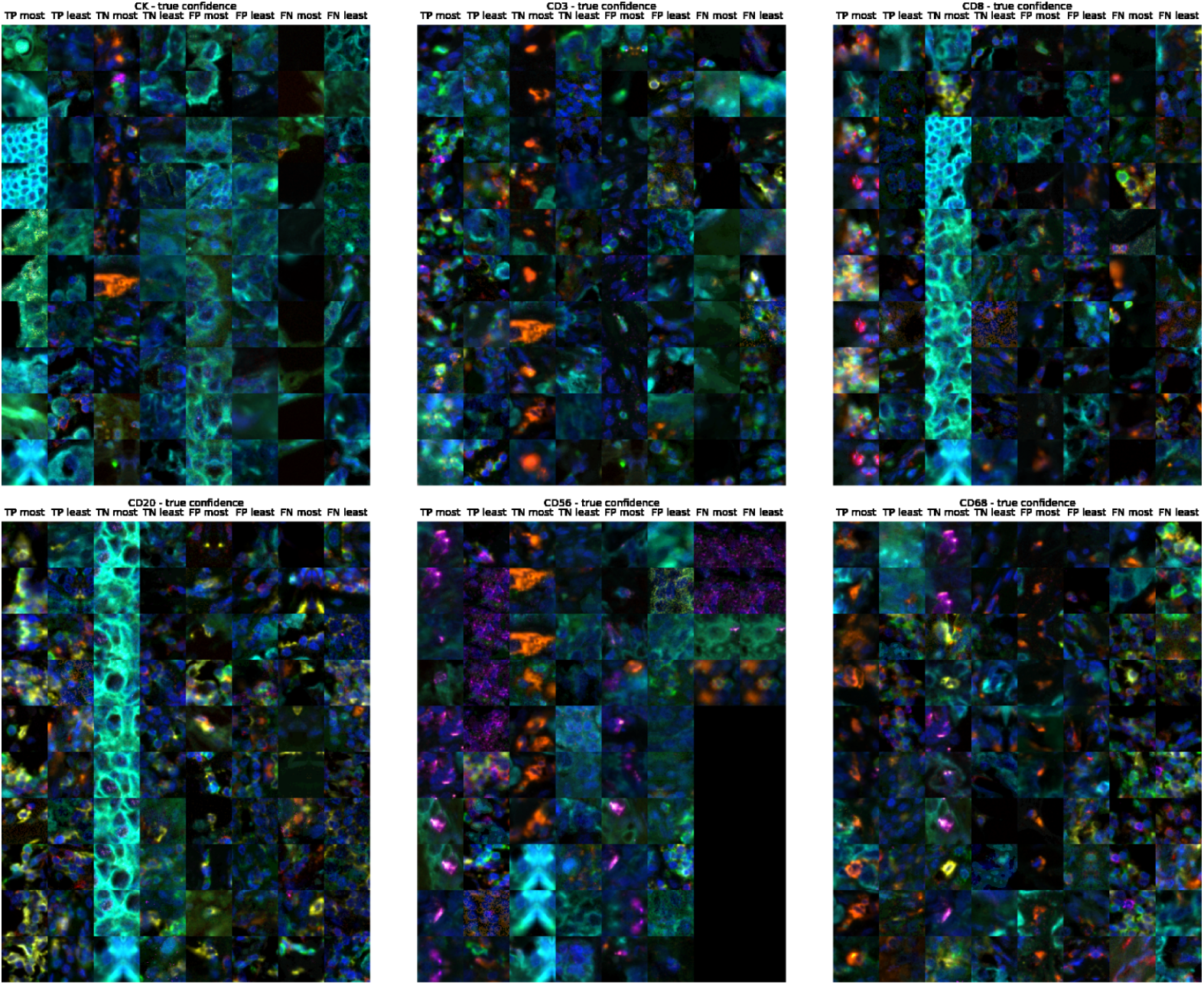
Example cell patch images of high/low confidence for TNBC1-MxIF8. Images per phenotype with the most and least confident predictions per classification outcome for the TNBC1-MxIF8 dataset with TME-Analyzer annotations. For each cell phenotype, a random sample of 10 cells drawn from the 100 most and least confident predictions is shown. Note that for CD56 only four false negative (FN) predictions were recorded.

**Fig. S16.**
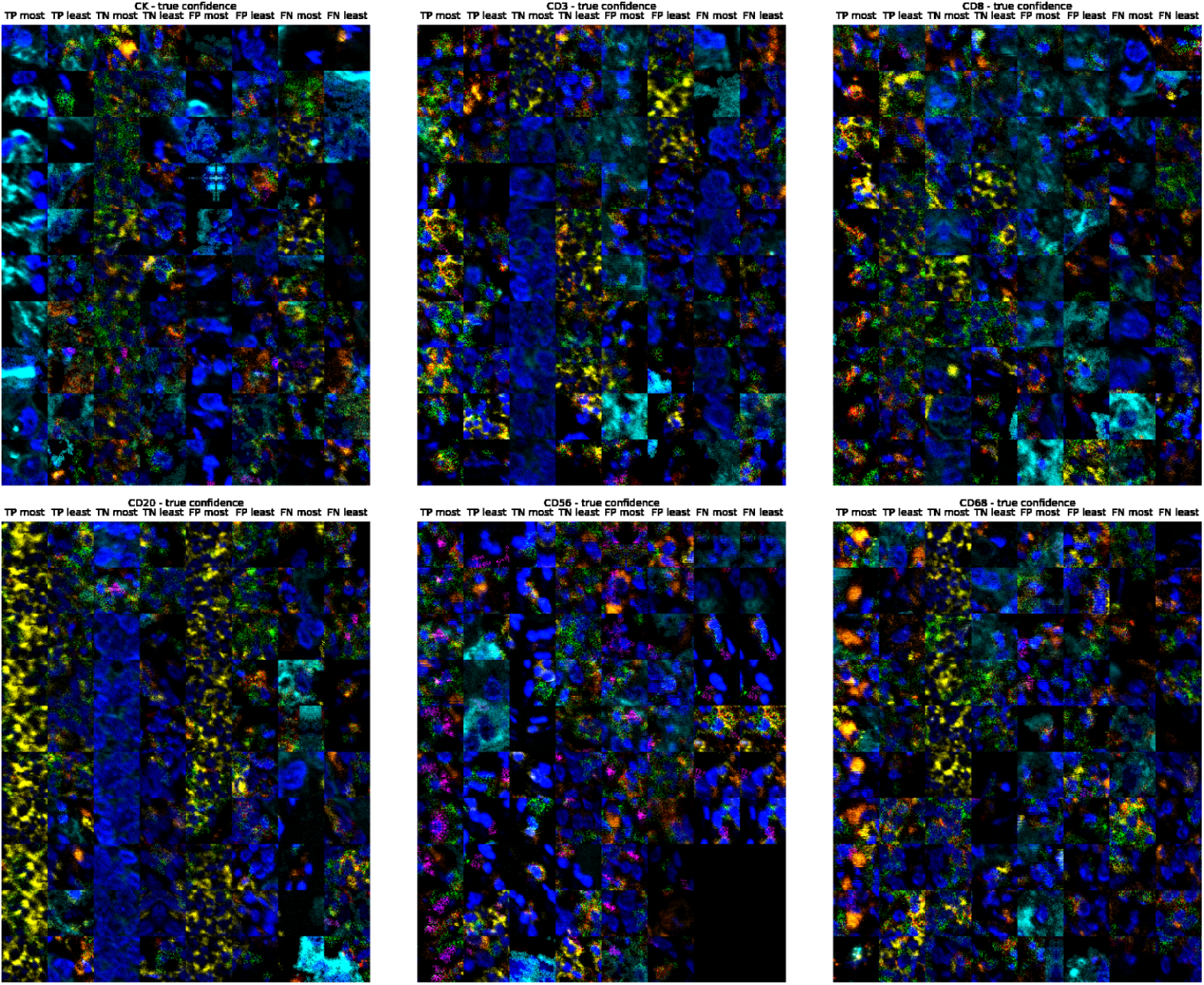
Example cell patch images of high/low confidence for TNBC2-MIBI8. Images per phenotype for the most and least confident predictions per classification outcome for the TNBC2-MIBI8 dataset with TME-Analyzer annotations. For each cell phenotype, a random sample of 10 cells drawn from the 100 most and least confident predictions is shown. Note that for CD56, only seven false negative (FN) predictions were recorded.

## Supplementary Tables

**Table S1.** Overview of patient data in the TNBC1-MxIF8 training set. The rightmost column specifies the mean of median (MoM) area under the precision-recall curve (PR-AUC) score obtained when training a classifier on data from only that patient (see **Fig. E5a**). Note that cell counts are based on the TME-Analyzer labels (TME-A ML6), which are multi-label, and the counts for individual class labels therefore do not necessarily add to the total number of cells.

| Patient ID | Cells total | CK+ | CD3+ | CD8+ | CD20+ | CD56+ | CD68+ | other cells | MoM (PR-AUC) |
| --- | --- | --- | --- | --- | --- | --- | --- | --- | --- |
| 1 | 26573 | 11071 | 477 | 273 | 13 | 11 | 45 | 14917 | 0.578099 |
| 2 | 46799 | 7042 | 3943 | 10121 | 8391 | 230 | 0 | 22484 | 0.524986 |
| 3 | 52719 | 10158 | 5756 | 9250 | 4804 | 7 | 656 | 29347 | 0.681697 |
| 4 | 31429 | 10059 | 593 | 1423 | 1775 | 196 | 0 | 18016 | 0.611027 |
| 5 | 36354 | 10069 | 1998 | 566 | 4141 | 26 | 60 | 20822 | 0.680865 |
| 6 | 26039 | 10738 | 195 | 81 | 8 | 1 | 10 | 15163 | 0.488509 |
| 7 | 66747 | 17104 | 1154 | 11013 | 11147 | 226 | 431 | 31453 | 0.708225 |
| 8 | 13240 | 5527 | 67 | 197 | 185 | 1 | 7 | 7387 | 0.442039 |
| 9 | 36731 | 9456 | 1163 | 6958 | 3795 | 7 | 844 | 16955 | 0.692290 |
| 10 | 63923 | 16298 | 10857 | 11768 | 7640 | 276 | 1984 | 27337 | 0.561818 |
| 11 | 6622 | 1955 | 224 | 13 | 10 | 2 | 15 | 4458 | 0.480732 |
| 12 | 13151 | 3421 | 1704 | 1213 | 320 | 11 | 186 | 7392 | 0.635892 |
| 13 | 28922 | 10807 | 388 | 954 | 23 | 1 | 2132 | 15859 | 0.607508 |
| 14 | 68345 | 17718 | 9831 | 8804 | 3607 | 381 | 5390 | 31315 | 0.680441 |
| 15 | 39085 | 14913 | 1344 | 605 | 266 | 6 | 100 | 22683 | 0.706655 |
| 16 | 50236 | 13951 | 2024 | 1711 | 600 | 514 | 677 | 32653 | 0.689051 |
| 17 | 27753 | 13096 | 309 | 476 | 35 | 851 | 0 | 13874 | 0.465894 |
| 18 | 37193 | 9794 | 1793 | 3390 | 1183 | 0 | 266 | 22571 | 0.578672 |
| 19 | 43491 | 9923 | 8572 | 10003 | 7472 | 261 | 0 | 17894 | 0.440875 |
| 20 | 41584 | 17499 | 1384 | 454 | 648 | 39 | 1246 | 21273 | 0.738653 |
| 21 | 77416 | 13275 | 7324 | 5703 | 3543 | 8 | 211 | 50726 | 0.712518 |
| 22 | 22102 | 7236 | 13 | 48 | 1 | 3 | 0 | 14806 | 0.380769 |
| 23 | 23880 | 8919 | 746 | 1334 | 1013 | 1 | 123 | 12410 | 0.559929 |
| 24 | 32084 | 9812 | 779 | 295 | 2 | 4 | 86 | 21421 | 0.541982 |
| 25 | 32588 | 13928 | 285 | 953 | 352 | 19 | 117 | 17282 | 0.619145 |
| 26 | 30219 | 4865 | 5829 | 3150 | 1400 | 7 | 140 | 17757 | 0.600300 |
| 27 | 18896 | 10782 | 155 | 560 | 0 | 18 | 901 | 7163 | 0.134859 |
| 28 | 37521 | 12510 | 6490 | 3278 | 2769 | 62 | 643 | 16722 | 0.542952 |
| 29 | 26502 | 10233 | 148 | 405 | 186 | 10 | 0 | 15692 | 0.474364 |
| 30 | 25141 | 7608 | 1062 | 478 | 161 | 95 | 1230 | 15099 | 0.631772 |
| 31 | 25541 | 8940 | 682 | 1365 | 4488 | 98 | 266 | 12531 | 0.512999 |
| 32 | 30974 | 8868 | 2112 | 666 | 594 | 48 | 3 | 19753 | 0.428093 |
| 33 | 42357 | 12753 | 5003 | 3589 | 1486 | 571 | 519 | 22321 | 0.779756 |
| 34 | 42212 | 15058 | 1573 | 614 | 684 | 13 | 537 | 24584 | 0.695987 |
| 35 | 60662 | 16857 | 4306 | 10525 | 4214 | 83 | 3770 | 30054 | 0.605924 |
| 36 | 43426 | 19077 | 2333 | 1448 | 1028 | 30 | 350 | 20769 | 0.743523 |
| 37 | 38205 | 14675 | 127 | 1152 | 158 | 142 | 59 | 22162 | 0.693791 |
| 38 | 44118 | 13573 | 1411 | 2648 | 1053 | 27 | 1369 | 25557 | 0.721065 |
| 39 | 32059 | 8882 | 295 | 2753 | 1733 | 48 | 744 | 18658 | 0.616728 |
| 40 | 16995 | 5930 | 70 | 482 | 429 | 3 | 0 | 10172 | 0.457956 |
| 41 | 40337 | 12172 | 6325 | 9217 | 5313 | 249 | 4118 | 12655 | 0.546611 |
| 42 | 45027 | 9517 | 5620 | 7751 | 4224 | 44 | 751 | 22643 | 0.641491 |
| 43 | 57606 | 19299 | 4078 | 11089 | 2039 | 28 | 4943 | 26500 | 0.657826 |
| 44 | 39783 | 14768 | 1906 | 1554 | 458 | 23 | 888 | 21368 | 0.719910 |
| 45 | 11836 | 5152 | 215 | 102 | 4 | 8 | 36 | 6495 | 0.446772 |
| 46 | 23814 | 6170 | 846 | 5799 | 1002 | 1 | 1545 | 10471 | 0.583151 |
| 47 | 53859 | 19000 | 2527 | 1039 | 691 | 8 | 2740 | 29650 | 0.726122 |
| Total | 1732096 | 530458 | 116036 | 157270 | 95088 | 4698 | 40138 | 919274 |  |

